# *APETALA2* controls seed growth by regulating outer integument mechanical identity during seed coat differentiation

**DOI:** 10.64898/2025.12.16.694797

**Authors:** Camille Bied, Runjue Yao, Audrey Creff, Luka Lelas, Debra David, Amélie Bauer, Sophy Boeuf, Guillaume Cerruti, Jeanne Braat, Alexandra Launay-Avon, Christine Paysant-Le-Roux, Frances Clark, Adrienne H. K. Roeder, Aline Voxeur, Gwyneth Ingram, John Golz, Benoit Landrein

## Abstract

Organ morphogenesis requires differentiating cells to acquire mechanical properties that precisely regulate growth. In *Arabidopsis thaliana*, seed size and shape are largely controlled by the outer integument of the seed coat, whose two layers develop distinct mechanical behaviors critical for growth regulation. Here, we show that the cell identity factor *APETALA2* (AP2) both promotes and restricts seed growth by coordinating the post-fertilization differentiation of these layers into distinct seed coat tissues. AP2 drives layer-specific remodeling of cell wall composition, notably affecting pectin and xyloglucan, thereby establishing contrasting mechanical properties between the two layers. Unexpectedly, these changes in wall composition and mechanics do not alter mechanosensitive responses, revealing a surprising uncoupling between cell wall remodeling and mechanotransduction. Our findings identify AP2 as a key coordinator of tissue-specific mechanical differentiation during organ morphogenesis.

## Introduction

The size and shape of organs arise from complex mechanical interactions among the various cells and tissues that compose them. These interactions generate internal forces that are perceived by cells and capable of influencing key processes involved in growth and patterning^1^. In plants, many organs grow as a result of inner tissues exerting outward pressure while being constrained by the mechanical properties of an outer layer put under tension, which is often the epidermis^2–4^. Cell walls in the epidermis exhibit unique mechanical properties that enable them to withstand the tensile forces generated by internal tissue pressure, while remaining sufficiently extensible to permit growth^2,5^. The epidermis also shows distinct mechanosensitive responses that influence these mechanical properties, thus contributing to organ morphogenesis^6–10^. Despite these insights, the molecular mechanisms that confer specific biomechanical properties to epidermal layers remain poorly understood, partly because mutants affecting epidermal identity are often lethal^11,12^.

Arabidopsis seeds provide an appealing model system to study the mechanical control of plant organ growth, as they comprise three genetically and physically distinct compartments: the embryo, the endosperm, and the seed coat—nested within one another like Russian dolls^13^. Seed growth, which occurs within a week after fertilization, emerges from mechanical interactions between two of these compartments, the endosperm and the seed coat^14^. Consistent with this view, our recent work has shown that seed size and shape are controlled by two distinct mechanosensitive responses triggered in the two layers of the seed coat outer integument in response to tension generated by endosperm expansion^15–17^. At early stages of seed development, microtubule responses to mechanical cues guide cellulose deposition according to shape-driven stresses in the outermost layer of the outer integument (the abaxial epidermis, ab-oi), thereby promoting seed elongation^17^. At later stages, mechanosensitive stiffening of the inner periclinal wall of the second layer of the outer integument (the adaxial epidermis, ad-oi), marked by the accumulation of demethylesterified pectin, restricts seed growth and sets final seed size^16^. Together, these studies support the idea that each layer of the seed coat outer integument possesses a distinct *mechanical identity* that endows its cell walls with specific mechanical properties essential for seed growth control, while also enabling tissue-specific responses to mechanical forces^14^.

Both layers of the outer integument are epidermal in origin, suggesting that their mechanical identities may not solely depend on epidermal factors^18^. In contrast, several studies have shown that the transcription factor APETALA2 (AP2), a well-known regulator of floral organ identity^19^, promotes outer integument differentiation, which affects final seed size, mass and yield^13,20,21^. Here, we investigated whether AP2 could control seed growth by modulating both the mechanical properties and the cellular responses to forces of each outer integument layer during seed development. Here, we show that APETALA2 (AP2) orchestrates the post-fertilization differentiation of each outer integument layer into a distinct cell type. AP2 directs layer-specific remodeling of cell wall composition, notably involving pectin and xyloglucan, which is essential for preserving cell integrity during growth and for controlling seed size and shape. Strikingly, despite these substantial changes in wall composition and mechanical properties, the layer-specific mechanosensitive responses that regulate growth remain unaffected, revealing an unexpected uncoupling between cell identity–driven wall remodeling and mechanotransduction. Together, our findings demonstrate how a transcriptional regulator of cell identity controls organ growth by selectively tuning the mechanical properties of individual cell layers, without altering their intrinsic mechanosensitive behavior.

## Results

### Antagonistic effects of AP2 on seed growth control

To investigate how *AP2* regulates seed growth through modulation of the outer integument properties, we first characterized the EMS-induced ap2*-6* mutant^22^ crossed with Wild Type (WT, Col-0 ecotype) pollen, ensuring that the *ap2* mutation was homozygous only in maternal tissues^13,20^. Similar to other *ap2* alleles^23^, *ap2-6* seeds lacked the characteristic differentiation features observed in WT seeds. In *ap2-6* seeds, the abaxial outer integument (ab-oi, also known as oi2) fails to form mucilage secretory cells (MSCs), while the adaxial layer (ad-oi, also known as oi1) showed defective differentiation, lacking starch granule accumulation at early stages and inner periclinal wall thickening (wall 3) at later stages (**Figure 1A**). To assess whether *AP2* expression coincides with these defects, we examined a *pAP2::AP2-VENUS* translational fusion, which complements the *ap2-12* null allele^24^. Confocal imaging revealed *AP2* expression in all cells of the outer integument in both mature ovules and developing seeds throughout the seed growth phase (**Figure 1B**), consistent with a role for this transcription factor in directing the differentiation of each outer integument layer during both ovule and seed development.

**Figure 1.**
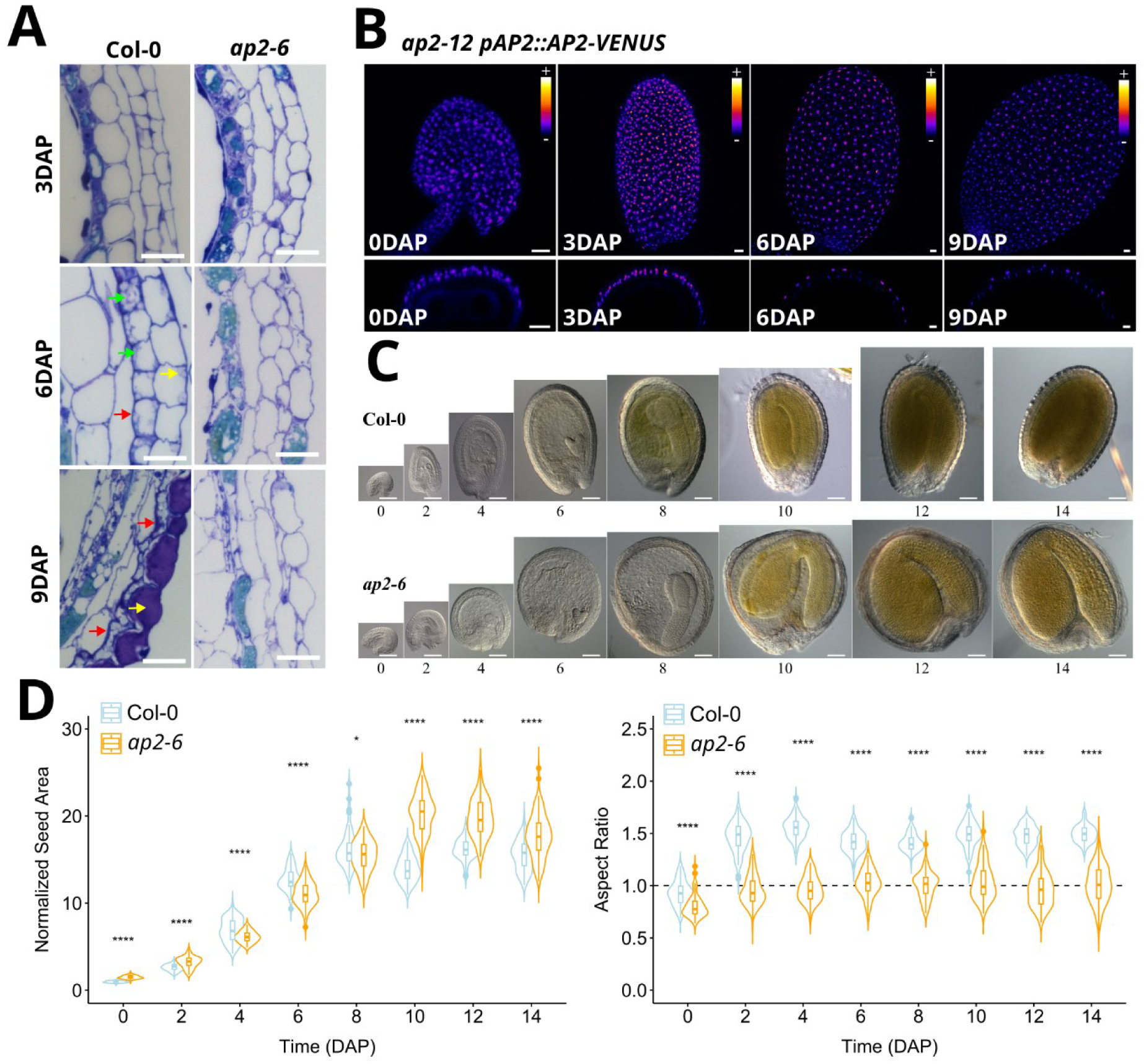
The transcription factor AP2 has two antagonistic effects on seed growth. **A.** Toluidine blue sections of WT and *ap2-6* seeds (crossed with WT pollen) showing the defects of outer integument differentiation of *ap2* mutants. Note the thickening of the inner-facing wall (red arrows) and the presence of starch granules (green arrows) in the ad-oi layer, and the progressive differentiation of the ab-oi layer into mucilage-producing cells (yellow arrow). Scale bars, 20µm. DAP: Days after pollination. **B.** Expression of *pAP2::AP2-VENUS* (in a *ap2-12* mutant background) in mature ovules and in developing seeds. Scale bar, 20 µm, *VENUS* intensity is shown using the Fire lookup table in ImageJ, n = 24 – 40 seeds from 5 to 6 plants from 2 independent experiments. **(C and D**) Representative pictures (C) and quantification (D) of the growth dynamics of WT and *ap2-6* mutant seeds (crossed with WT pollen). Scale bar, 20µm, n = 63 – 119 seeds from 3 plants per day per genotype, one experiment. Data were compared using bilateral Student’s tests.

To determine how *AP2*-mediated differentiation of the outer integument affects seed growth, we compared growth dynamics of WT and *ap2-6* seeds fertilized with WT pollen during the first 12 Days After Pollination (DAP). Consistent with *AP2* activity in the ovule integument, *ap2-6* ovules were larger than WT prior to fertilization (**Figure 1C–D**, **Figure S1A**). After fertilization, however, *ap2-6* seeds grew more slowly, resulting in smaller seeds relative to WT at 4–6 DAP. At later stages (>6 DAP), *ap2-6* seed growth decelerated less sharply, ultimately producing larger seeds than WT. Morphometric analyses further revealed that *ap2-6* ovules were slightly more elongated than WT but failed to undergo anisotropic expansion during the first 3 DAP, when WT seeds normally exhibit rapid directional growth^17^ (**Figure 1C–D**, **Figure S1A**). Consequently, *ap2-6* seeds developed a rounder shape. These results show that AP2 activity in the seed coat affects ovule development, and plays two opposing roles in seed growth: it promotes early anisotropic expansion but later limits isotropic growth.

To validate these findings, we characterized the phenotype of additional *ap2* mutant alleles of varying strength, as determined by their floral phenotypes^25,26^. The weak EMS-induced *ap2-5* allele, which causes only partial homeotic conversion of petals into stamens in flowers, showed elongated ovules similar to *ap2-6* but, crossed with WT pollen, produced seeds that were either slightly smaller or comparable in size to WT, and only marginally less elongated (**Figure S1B–D**). In contrast, the EMS-induced *ap2-7* and the T-DNA insertion *ap2-12* alleles, which exhibit more frequent homeotic conversion of sepals and petals into carpels and stamens, displayed growth defects similar to those observed in *ap2-6* when crossed with WT pollen, despite differences in the severity of their floral phenotypes^26^ (**Figure. S1B-D**).

### AP2-dependent specification of seed coat outer integument identity

To elucidate the molecular mechanisms through which *AP2* regulates outer integument differentiation, we performed bulk RNA sequencing on WT and *ap2-6* (crossed with WT pollen) seeds at two developmental stages (2 and 5 DAP). We identified 1,300 and 1,155 differentially expressed genes (DEGs) at 2 and 5 DAP, respectively (|log₂FC| ≥ 2, *p* ≤ 0.05), among which 58 – 60% were downregulated, with a ∼14% overlap between stages (**Figure 2A**, **Table S1, Figure S2A** for a lower cutoff). Gene ontology (GO) analysis revealed enrichment in categories related to “Cell Wall” (primary wall at 2DAP; secondary wall at 5 DAP), and “Glucosinolate Metabolism” at both stages, as well as “Response to Stress” at 2DAP (**Figure 2B, Figure S2B**) and “Transcriptional Regulation” at 5DAP, consistent with *ap2* defects in seed coat growth and differentiation.

**Figure 2.**
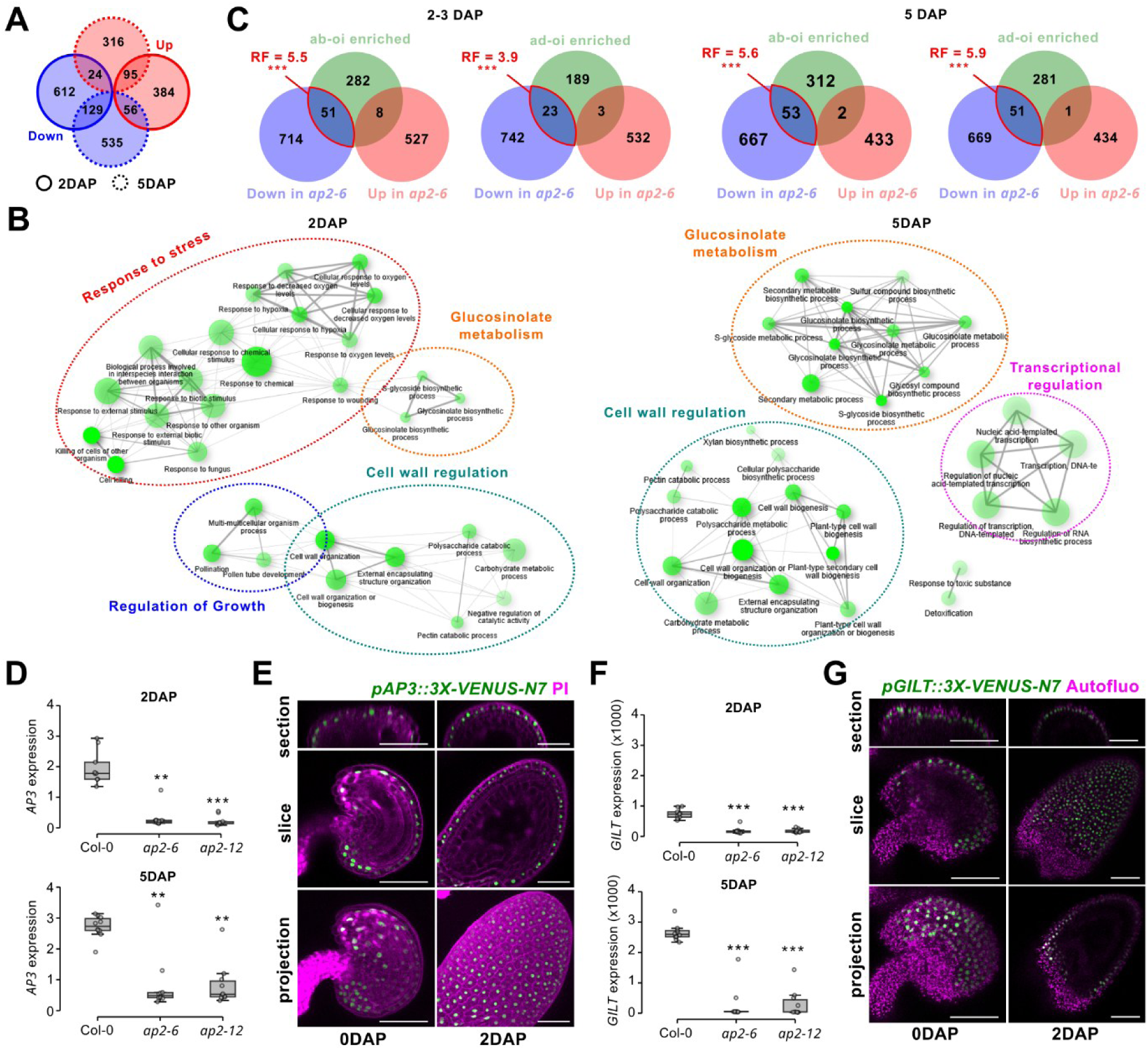
AP2-dependent control of outer integument identity. **A.** Number and overlap of the DEGs down or up-regulated at 2 and 5DAP in *ap2-6* seeds (crossed with WT pollen) compared to the WT (|log₂FC| ≥ 2, *p* ≤ 0.05). **B.** Network representation of the 30 most enriched Gene Ontologies (GOs) in *ap2*-6 DEGs at 2 and 5DAP (|log₂FC| ≥ 2, *p* ≤ 0.05). In this representation, obtained with ShinyGO, two GOs are connected if they share at least 30% of genes, thicker lines represent more overlapped genes, while larger and brighter spots represent more connected GOs. **B.** Overlap between the genes enriched in the ab-oi or ad-oi layer at 3 and 5 DAP (|log₂FC| ≥ 2, *p* ≤ 0.05, data from^30^) and the genes under and over-expressed in *ap2-6* at 2 and 5 DAP (|log₂FC| ≥ 2, *p* ≤ 0.05), respectively. Each representation factor (RF) and its significance was calculated using a Fisher’s Exact Test for Count Data. **C, E** Relative expression of *AP3* (C) and *GILT* (E) at 2 and 5 DAP in the WT and in the *ap2-6* and *ap2-12* mutants (crossed with WT pollen); n = 10 replicates from 2 independent experiments. Statistical differences established using the Kruskal-Wallis test followed by Dunn’s Post-hoc test. **D, F** Representative pictures showing the expression pattern of *AP3* (D, imaged using a *pAP3::3X-VENUS-N7* reporter) and *GILT* (F, imaged using a *pGILT::3X-VENUS-N7* reporter) in unfertilized ovules (0DAP) and in developing seeds (2DAP). Scale bars, 20 µm; *AP3*: n = 36 – 46 seeds per timepoint from 7 – 10 plants from 2 experiments; n= *GILT:* n = 26 seeds per timepoint from 6 plants from 2 independent experiments

Because seed coat development can non-autonomously affect zygotic tissues^27,28^, we next examined whether *ap2-6* DEGs originated from the seed coat or from other seed compartments. Using a published microarray dataset^29^, we found that only a subset of *ap2-6* DEGs corresponded to seed coat–enriched genes, while many were predominantly expressed in the endosperm and, to a lesser extent, in the embryo (**Figure S2C**). To better resolve seed coat-specific expression, we thus compared our RNAseq data with two recent single-nucleus RNA-seq (snRNAseq) datasets of developing seeds^30,31^. We found that there was significant overlap between the genes normally enriched in each outer integument layer at 3 and 5 DAP and the genes that were downregulated in *ap2-6* at 2 and 5 DAP, respectively (representation factor 3.9 – 5.9, p<0.001, **Figure 2C**, **Tables S2–S5,** see **Figure S2D** for a lower cutoff). These results show that *AP2* is required for the proper activation of at least part of the tissue-specific transcriptional program of each outer integument layer. To validate these transcriptomic results, we performed quantitative PCR (qPCR) and live imaging of fluorescent reporters. We found that marker genes such *AP3* and *GILT*, whose expression is, according to published snRNAseq datasets, enriched in the ab-oi and ad-oi layers, respectively^30,31^, showed layer-specific expression in both WT ovules and developing seeds, and were downregulated at 2 and 5 DAP in *ap2-6* and *ap2-12* mutant alleles (**Figure 2C–F**).

We next examined the expression of transcription factors (TFs) that could act in the control of outer integument differentiation downstream of *AP2*. We found that the expression of several TFs specific to the ad-oi layer^30^, putative direct AP2 targets according to a published a ChIP-RAseq dataset (Chromatin Immunoprecipitation)^24,25^, was strongly downregulated in *ap2* seeds, making them strong candidates for mediating ad-oi differentiation downstream of AP2 (**Tables S6–S8**, **Figure S3A**). In the ab-oi layer, expression of several genes involved in epidermal specification and/or differentiation^32^, such as the HD-ZIP IV TF-encoding genes *GL2*, and *HDG2*, was downregulated in *ap2-6* (**Tables S9–S10, Figure S3C**). We therefore asked specifically how epidermal identity is influenced by AP2 activity. We examined the expression of the epidermal regulators *ATML1*^33^, *PDF2*^33^, and *GL2*^34^, and of the epidermal marker *PDF1*^35^, using fluorescent reporters. In mature ovules, all reporters were active in every integument layer, consistent with the epidermal (L1) origin of the ovule integuments^18^ (**Figure 3A, Figure S3B**). During seed development, all reporters became restricted to the ab-oi layer and, to a lesser extent, to the ad-oi layer for *ATML1* and *PDF2*, while weak expression was also detected in the endothelium (**Figure 3A**). In *ap2-6* mutants, expression patterns of all reporters resembled those of WT ovules but diverged after fertilization (2 DAP). *ATML1* and *PDF2* expression remained largely unaffected in all seed coat integuments (**Figure S3B**), whereas GL2 and *PDF1* expression in the ab-oi layer was strongly affected, becoming patchy for *GL2* and very weak for *PDF1*, while remaining weak and homogeneous in the endothelium (**Figure 3B–C**). These data indicate that AP2 is partly required for the correct acquisition of ab-oi seed coat identity after fertilization, as characterized by *GL2* and *PDF1* expression.

**Figure 3.**
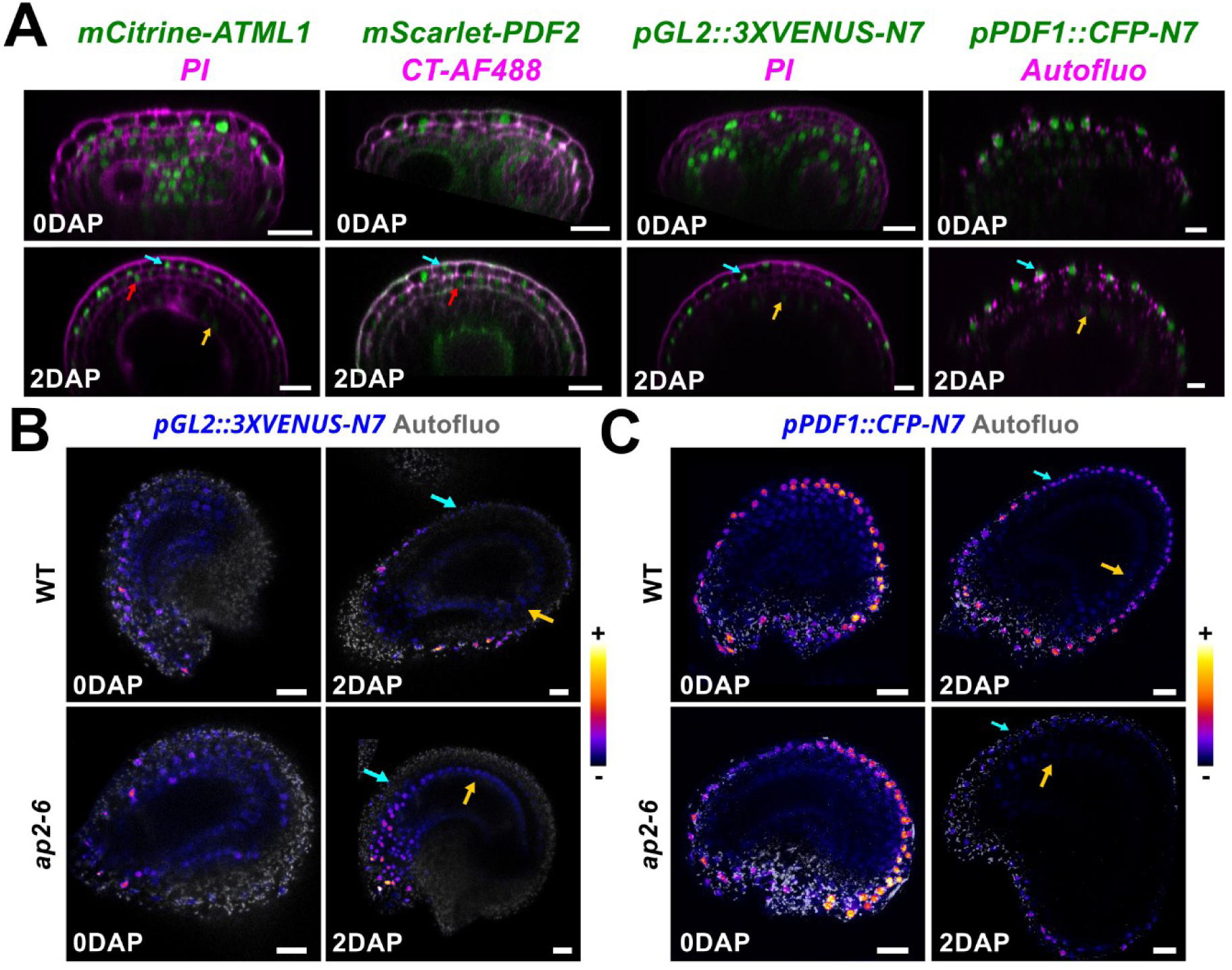
AP2-dependent expression of genes associated with epidermal fate after fertilization. **A.** Representative confocal section of unfertilized ovules (0DAP) and developing seeds (2DAP) expressing transcriptional and translational reporters for the epidermis-associated genes *ATML1*, *PDF2*, *GL2*, and *PDF1*. Scale bars, 10 µm. **B-C** Representative sections (5 slices z-projections) of mature ovules (0DAP) and developing seeds (2DAP) of WT and *ap2-6* seeds (crossed with WT pollen) expressing transcriptional reporters for *GL2* (B, *pGL2::3X-VENUS-N7*) and *PDF1* (C, *pPDF1::CFP-N7*). *GL2:* n=30 - 45 seeds from 6 - 7 plants, *PDF1*: n= 35 – 37 seeds from 7 – 8 plants, two independent experiments; Scale bars, 20 µm. Blue, red, and yellow arrows point at ab-oi, ad-oi and endothelium nuclei, respectively.

*GL2* expression in the ab-oi layer is thought to depend on activation of the MYB–bHLH–WD repeat (MBW) pathway, which drives the late differentiation of the ab-oi layer into MSCs^36–39^, a process impaired in *ap2* mutants (**Figure 1A**). We noticed that the expression of several MBW pathway components, such as *MYB5*, *MYBL2*, *TTG2*, *GL2*, and *HDG2,* was downregulated in *ap2* mutant alleles at 2 and 5 DAP (**Figure S3B**). We thus hypothesize that this pathway, which controls epidermal differentiation in other contexts^40^, may be activated upon fertilization to drive ab-oi layer specialization, promoting early changes in cell properties that are important for growth control (in agreement with the growth defects of *ttg2* mutants^41,42^), and subsequent differentiation into MSCs (**Figure S3D**).

Together, these findings reveal that seed coat identity specification involves a two-step process; an initial, pre-fertilization phase marked by layer-specific *AP3* and *GILT* expression is followed by post-fertilization refinement revealed by epidermal reporter activation in the ab-oi layer, and that depends in part on AP2 activity. We next examined how the defects in outer integument identity specification of *ap2* mutants could contribute to the seed growth defects observed after fertilization.

### AP2-dependent regulation of wall properties after fertilization

We previously showed that outer integument cells undergo progressive changes in wall composition that are essential for controlling seed size and shape^16,17^. In primary walls, such as those of the seed coat during growth, cellulose fibers are the main load-bearing elements, but wall stiffness and extensibility also depend on the pectin matrix and hemicelluloses connecting cellulose fibers^43^. In our transcriptomic analysis, we found that many cell wall regulators predicted to be expressed in the WT outer integument are downregulated in *ap2-6* seeds at 2 or/and 5DAP. This notably included pectin modifying enzymes such as *PME1, PME6, PME44* and *PAE12*; and hemicellulose-modifying enzymes such as *IRX8*, *XTH7*, *XTH14* and *XTH33* (**Table S11-S14**). We thus studied the impact of AP2 activity on pectin and hemicellulose content of seed coat walls.

Enzymatic fingerprinting of whole seeds with an endopolygalacturonase, which allows the characterization of fine structure of low- and non-methylesterified pectins, revealed developmental and genotype-specific differences in the composition in homogalacturonans (HGs), the most abundant pectic polysaccharide in primary walls^44^ (**Figure 4A**; **Figure S4A**). In *ap2-6*, digestible HGs were initially slightly less methylesterified compared to WT. During seed development, digestible HGs became progressively less methylesterified, both in WT and *ap2-6*. By contrast, digestible, non methylesterified and acetylated HGs accumulated over time in the WT, but not in *ap2.6* seeds. Immunolabeling with pectin-specific antibodies confirmed major differences in HG methylesterification patterning between outer integument walls in WT seeds (pectin acetylation could not be probed because of a lack of specific antibody). Methylesterified pectins (LM20) accumulated in wall 1, while partially demethylesterified pectins (LM19, JIM5) localized mainly to walls 2 and 3 and increased over time (**Figure 4B**; **Figure S4B**). Unesterified, calcium-crosslinked pectins (2F4) appeared specifically in wall 3 at later stages. In *ap2-6* seeds, LM20 labelling in wall 1 and 2F4 labelling in wall 3 were lost, while LM19 and JIM5 signals were more uniform across the walls of both ab- and ad-oi layers. Analysis of additional *ap2* alleles with LM20, LM19 and JIM5 confirmed these trends: *ap2-5* resembled WT, whereas *ap2-7* and *ap2-12* showed *ap2-6*-like patterns (**Figure S4B-C**), consistent with the severity of their growth defects (**Figure S1C-D**). Together, these results indicate that AP2 is required for the proper spatial patterning and maturation of pectins in the seed coat.

**Figure 4.**
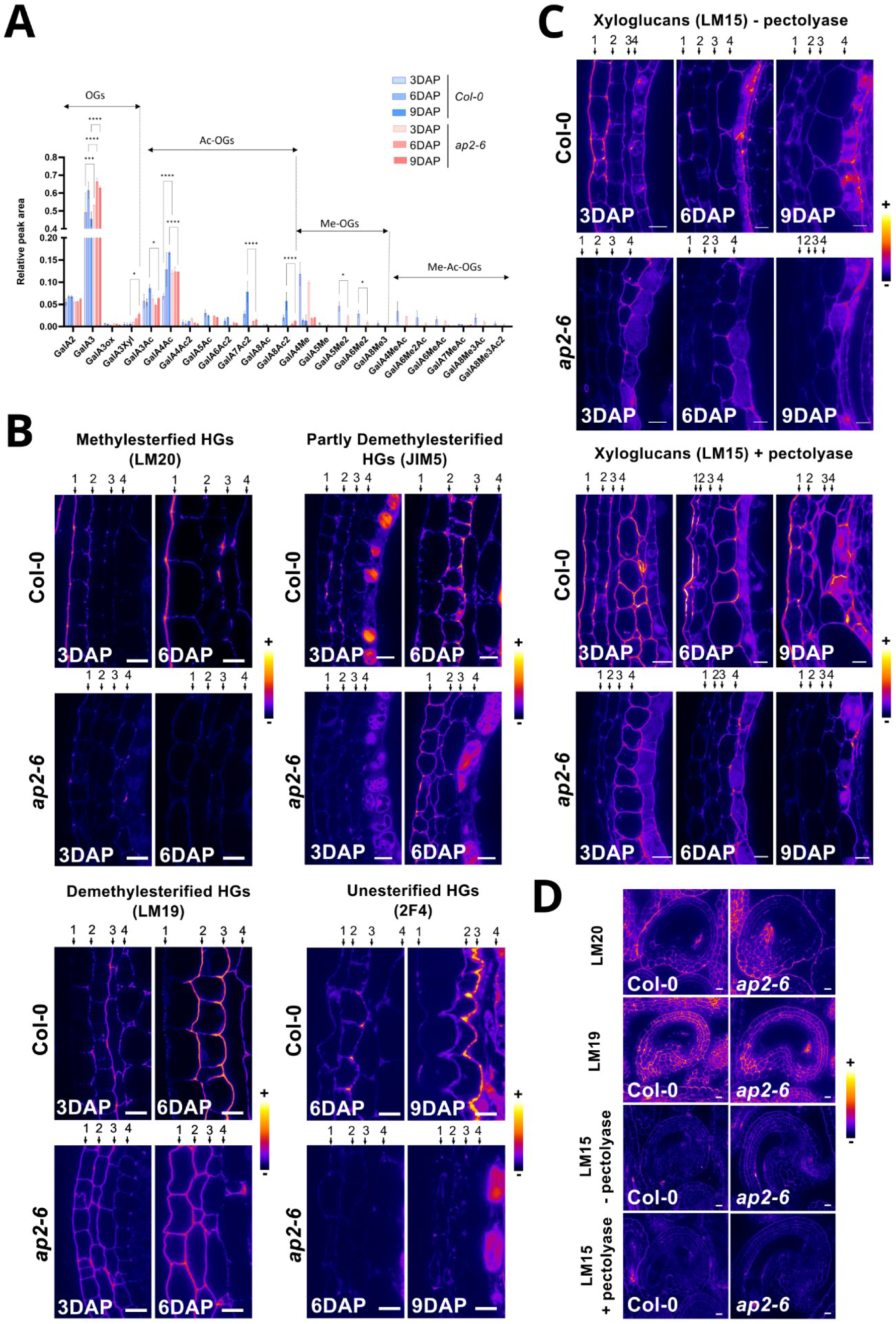
Post-fertilization defects in outer integument wall composition in *ap2-6* mutant. **A.** Relative level of the oligogalacturonides released following digestion of intact seeds at 3, 6 and 9 DPA with a polygalacturonase. One experimental repeat with 3 biological replicates, WT and *ap2-6* samples (crossed with WT pollen) were compared using bilateral Student’s tests. **B.** Representative signal of the LM20 (labelling methylesterified homogalacturonans, JIM5 and LM19 (labelling partially demethylesterified homogalacturonans) and of the 2F4 (labelling unesterified pectins) antibodies in WT and *ap2-6* (crossed with WT pollen) seeds at 3 and 6 or 6 and 9DAP. Scale bars, 10 µm; n = 9 – 18 seeds, two independent experiments. **C.** Representative signal of the LM15 antibody (labelling xyloglucans, without or with pectolyase treatments) in WT and *ap2-6* (crossed with WT pollen) seeds at 3, 6 and 9DAP. Scale bars, 10 µm; n = 9 – 11 seeds, two independent experiments. **D.** Representative signal of the LM20, LM19, and LM15 (without or with pectolyase treatment) antibodies in WT and *ap2-6* ovules before fertilization. Scale bars, 10 µm; n = 8 – 22 ovules, two independent experiments. The LM19 and LM20 staining of mature ovule in D were done on independent batches of plants from the ones of developing seeds in C, but not the LM15 staining that are shown with the same look up table in panels B and C for comparison.

We next analyzed xyloglucan, the main hemicellulose in primary cell walls^43^. Endocellulase fingerprinting released distinct oligosaccharides depending on developmental stage but showed minimal genotype-dependent differences across experimental replicates (**Figure S4D**). In contrast, LM15 immunolabeling revealed marked differences in xyloglucan distribution. In WT, LM15 signal was detected in all outer integument walls, particularly in wall 1. Signal was detected only after pectolyase treatment at 6 and 9 DAP, indicating increasing xyloglucan–pectin association over time^45^. In *ap2-6*, LM15 labeling was strongly reduced, even after pectolyase treatment (**Figure 4C**; **Figure S4B**). Thus, AP2 activity significantly affects both pectin and hemicellulose composition in outer integument cell walls, correlating with the reduced expression of pectin- and xyloglucan-associated enzymes in *ap2-6* seeds (**Table S11-S14**).

Because AP2 is expressed throughout ovule and seed development, we tested whether these wall defects also occur before fertilization. In WT, the patterns of HG methylesterification in mature ovules were different from those observed in developing seeds: LM20, which detects methylesterified pectins, was still enriched in wall 1 but LM19, detecting demethylesterified pectins, marked all seed coat walls (**Figure 4D**). LM15 signal was very weak in mature ovules compared to developing seeds, even with pectolyase treatment, supporting reduced xyloglucan levels. Strikingly, no marked differences in pectin (LM19, LM20) or xyloglucan (LM15) distribution were observed between WT and *ap2-6* ovules (**Figure 4D**; **Figure S4C**). These data show that AP2 activity is specifically required for wall remodeling associated with seed coat differentiation, consistent its function in specifying outer integument identity after fertilization.

To study if changes in wall composition affect wall structure, we first measured outer integument wall thickness. In WT seeds, wall 1 is known to thicken early, consistent with the load-bearing function of the ab-oi layer at early stages of seed development^17^. In *ap2-6*, however, we observed that wall 1 thickening was strongly reduced, but partially compensated by a thicker wall 2 (TEM, **Figure 5A**). At later stages, we also observed that wall 3 thickened in WT but not in *ap2-6* (Toluidine blue, **Figure 1A**). Together with the uniform LM19 and JIM5 labeling of *ap2-6* outer integument walls (**Figure 4C**), these data suggest that AP2 controls not only wall composition but also the cell polarization that allows targeted deposition of wall components and their regulators, enabling specific wall face to acquire specialized properties.

**Figure 5.**
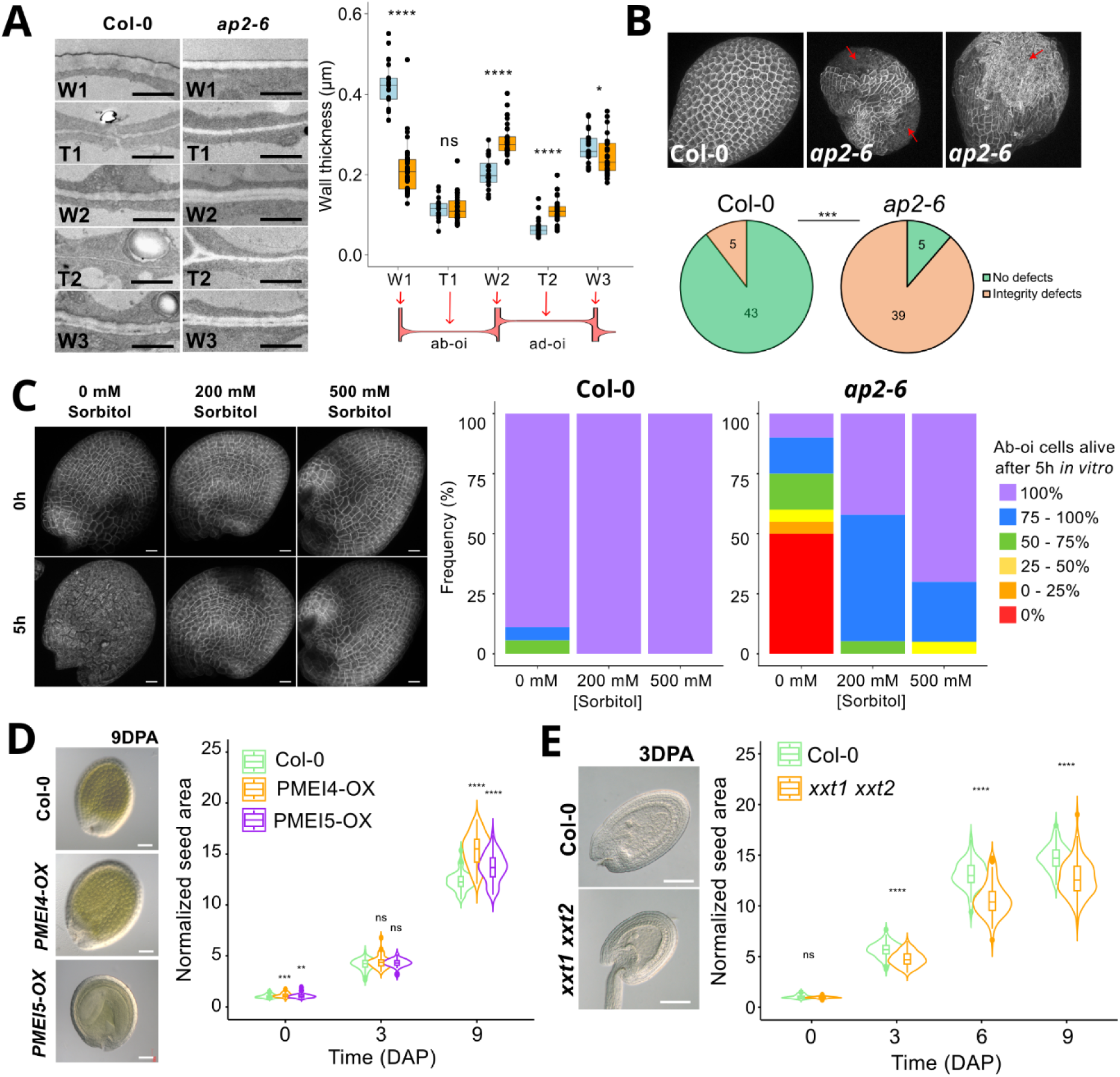
AP2-dependent control of wall properties and seed growth likely occurs through pectins and xyloglucans. **A.** Representative pictures and quantification of the thickness of outer integument cell walls in WT and *ap2-6* (crossed with WT pollen) seeds at 3DPA based on sections imaged by Transmission Electron Microscopy (TEM). Scale bars: 1 µm, 1 experimental repeat with 7-21 cells from 4-5 seeds per genotype. Data were compared using bilateral Student’s tests. **B.** Representative WT and *ap2-6* seeds (crossed with WT pollen) expressing the *LTi6b-GFP* membrane marker observed by confocal microscopy and showing the cell death phenotype of *ap2-6* seeds at 5DAP. Scale bars, 20 µm; n= 44 – 48 seeds from 3 independent experiments, frequencies were compared with a χ² test. **C.** Effect of medium osmolarity on ab-oi cell survival in WT and *ap2-6* seeds expressing a membrane marker at 2 DAP and cultivated *in vitro* for 5h; n= 18 – 20 seeds per condition, pool of two independent experiments; scale bars, 20 µm. **D.** Growth dynamics of WT, *PMEI4-OX* and *PMEI5-OX* seeds at 0, 3 and 9 DAP; n = 31 – 84 seeds from 3 siliques per day per genotype, one experiment. Data were compared using bilateral Student’s tests. **E.** Growth dynamics of WT and *xxt1 xxt2* seeds at 0, 3, 6 and 9 DAP; n = 42 – 124 seeds from 3 siliques per day per genotype, one experiment. Data were compared using bilateral Student’s tests.

We also attempted to examine the mechanical properties of *ap2* walls, but mechanical testing of *ap2-6* outer integument walls proved difficult due to their fragility, which prevented us from cultivating *ap2* seeds in vitro for extended periods. Consistent with this, we observed pronounced wall-integrity defects in the *ap2-6* mutant, leading to frequent death of ab-oi cells from 5 DAP onward (**Figure 5B**). To test whether these integrity defects result from *ap2* outer integument walls being unable to withstand pressure-induced tension, we cultivated isolated WT and *ap2-6* seeds at 2 DAP in media containing various concentrations of sorbitol. Despite some variation between experimental repeats, we found that the death of ab-oi cells normally observed in *ap2-6* seeds after 5 h of in vitro culture could be alleviated by increasing the osmolarity of the medium to decrease seed turgor (**Figure 5C**, **Figure S5A**). These findings support that AP2 activity is required to maintain outer integument wall integrity during growth.

To assess whether altered wall composition could contribute to the *ap2* growth phenotype, we next analyzed transgenic lines with modified pectin and xyloglucan content. Overexpression of *PECTIN METHYL ESTERASE INHIBITORS* (*p35S::PMEI4-GFP*^46^; *p35S::PMEI5*^47^) caused a modest increase in ovule and early seed size (3 DAP), and a stronger enlargement at late stages (9 DAP), particularly in *PMEI4*-OX (**Figure 5D, Figure S5B-C**). This indicates that pectin demethylesterification is principally required to restrict late seed growth, consistent with the idea that unesterified, calcium-crosslinked pectins in wall 3 stiffen the wall to limit expansion^16^, a process impaired in *ap2* mutants. To assess the role of hemicellulose, we analyzed the *xxt1 xxt2* double mutant, which lacks xyloglucans^48^. Early seed growth (0–3 DAP) was reduced in this double mutant, producing smaller, rounder seeds than WT, similar to early *ap2* developing seeds (**Figure 5E, Figure S5D-E**). This supports that xyloglucan contributes to early anisotropic seed expansion, likely by facilitating cellulose microfibril sliding in wall 1^49,50^; although pectins may also influence early growth, as suggested by altered LM19/LM20 patterns in *ap2* walls at early stages, and the reduced growth anisotropy observed in *PMEI5*-OX (**Figure S5B-C**). Together, our results support that AP2 modulates seed growth by regulating the post-fertilization composition of pectins and hemicelluloses in the outer integument walls. We next wondered whether AP2 might also be required for outer integument cells to exhibit specific mechanosensitive responses.

### AP2-independent tissue-specific mechanosensitive responses

Our previous work showed that the two layers of the outer integument exhibit distinct mechanosensitive behaviors that are crucial for controlling seed size and shape^14,16,17^. Given the role of *AP2* in seed coat differentiation and the associated wall defects of *ap2* mutant seeds, we asked whether AP2 activity is also required for tissue-specific mechanosensitive responses in the seed coat.

We first examined the ad-oi–specific gene *EUI-like p450 A1* (*ELA1)*, whose expression is induced in WT seeds upon mechanical stimulation^15,16^. The *pELA1::3X-VENUS* reporter showed similar expression patterns in the ad-oi layer of WT and *ap2-6* seeds before and after fertilization (**Figure 6A**), but *ELA1* transcript levels were reduced in the mutant at 5DAP, as determined by qPCR (**Figure S6A**). Moreover, *ELA1* expression could still be induced in *ap2-6* seeds at 2DAP by compressing fruits for 24h. These results indicate that *AP2* is not required for the mechanosensitive activation of this gene in the ad-oi layer (**Figure 6B**).

**Figure 6.**
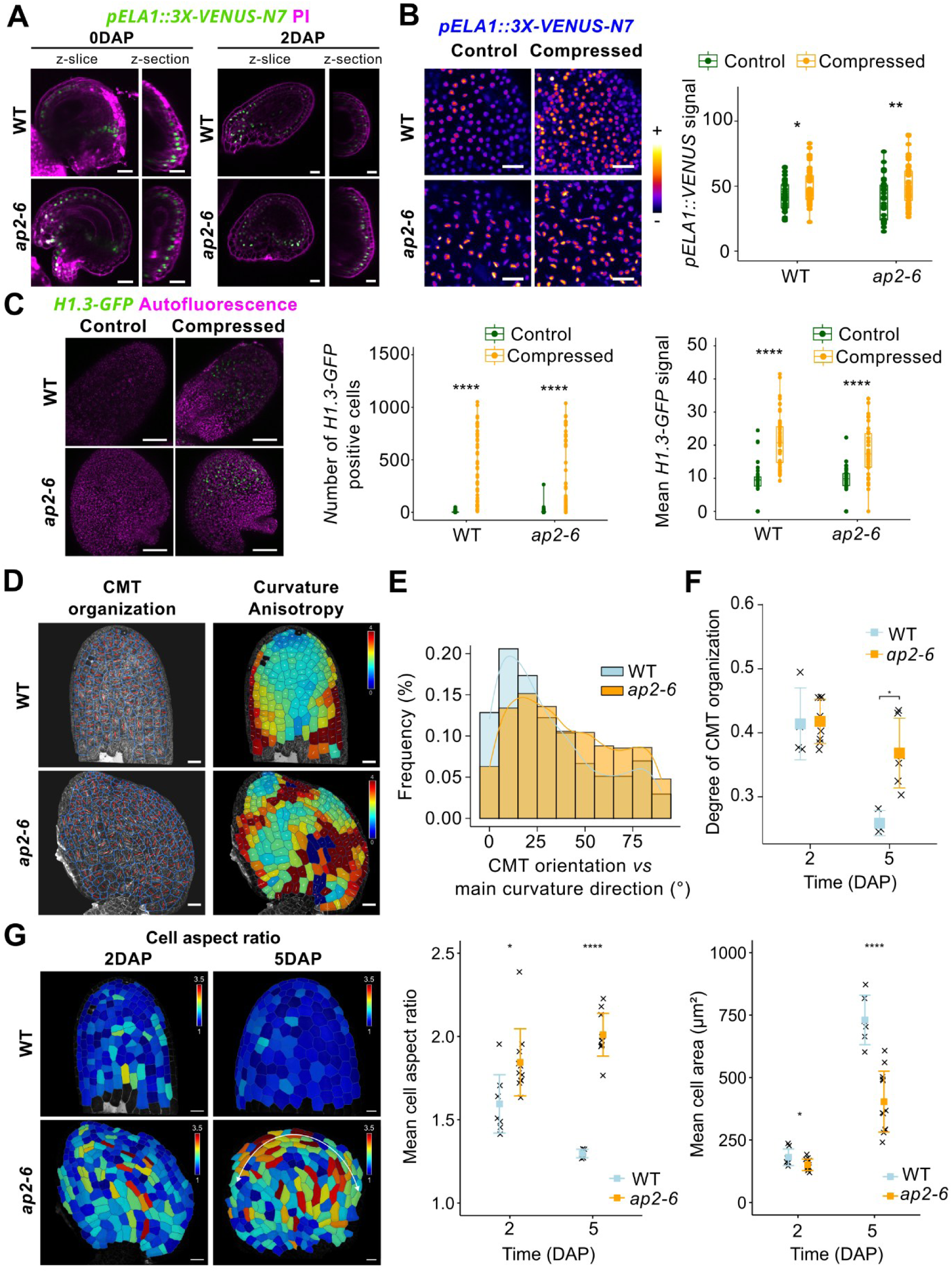
Mechanosensitive responses in *ap2-6* outer integument. **A.** Representative pictures showing the expression of *ELA1* (using the *pELA1::3X-VENUS-N7* reporter) in WT and *ap2-6* mature ovules (0DAP) and developing seeds (2DAP). Scale bars: 20µm; 2 DAP: n = 14 – 19 seeds from 3 plants, 5DAP: n = 26 – 35 seeds from 7 - 9 plants, 3 independent experiments. **B.** Effect of a 24h compression of the fruit with a microvice on *ELA1* expression in WT and *ap2-6* (crossed with WT pollen) mutant seeds assessed using the *pELA1::3X-VENUS-N7* reporter. Scale bars, 20 µm, n = 76 – 399 cells from 33 – 45 seeds from 7 – 10 plants per genotype per condition, 2 independent experiments. Data were compared using bilateral Student’s tests. **C.** Effect of a 24h compression of the fruits with a microvice on *H1.3* expression in WT and *ap2-6* mutant seeds assessed using the *pH1.3::H1.3-GFP* reporter. Scale bars, 20 µm, n = 0 – 1051 detected cells from 45 – 48 seeds from 10 plants per genotype per condition, 2 independent experiments. Data were compared using bilateral Student’s tests. **D.E** Representative pictures (D) and quantification of the correlation (E) between the orientation of the CMTs (imaged using the *p35S::MAP65-1-RFP* reporter) facing the outer-side of the ab-oi layer and the curvature anisotropy in WT and *ap2-6* (crossed with WT pollen) seeds at 2 DAP. Scale bars: 20 µm, n = 149 – 246 cells from 4 – 8 seeds per genotype, two independent experiments. For the CMT picture, in each cell highlighted in blue and the orientation of the red bars shows the mean orientation of the CMTs in each cell and their length, their degree of organization. In the heatmap of curvature anisotropy, the orientations of the two perpendicular white bars represent the axes of maximum and minimum curvature in each cell, and their lengths the degree of curvature in each of these two directions. **F.** Mean degree of organization of the CMTs in the ab-oi layer of WT and *ap2-6* (crossed with WT pollen) seeds at 2 and 5 DAP, n = 3 – 8 seeds, two independent experiments for 2DAP, one for 5DAP. Data were compared using bilateral Student’s tests. **G.** Heatmap representation and quantification of the shape (aspect ratio) and size (area) of the cells in the ab-oi layer of WT and *ap2-6* (crossed with WT pollen) seeds at 2 and 5 DAP. Scale bars: 20 µm, n = 80 – 403 cells from 6 to 12 seeds, 2 independent experiments. Data were compared using bilateral Student’s tests. In the heatmap of cell aspect ratio at 5DAP, the white line shows the elongation of the radial elongation of the cells at the margin of the seed in *ap2-6*.

We next analyzed the expression of the osmo- and mechano-sensitive histone variant *H1.3*^51–53^. Both RNA-seq and qPCR analyses revealed that *H1.3* expression was upregulated in *ap2* mutants at 2 DAP (**Table S1**, **Figure S6B**), consistent with the enrichment of the GO term “Response to Stress” among *ap2-6* DEGs (**Figure 2A**). Under control conditions, only a few nuclei expressed *H1.3-GFP* in both WT and *ap2-6* seeds, similar to the restricted expression pattern of this reporter in the shoot apical meristem in normal conditions^51^. However, strong *H1.3-GFP* induction was observed in both WT and *ap2-6* outer integument after compression (**Figure 6C, Figure S6C**), further confirming that AP2 is not required for outer integument cells to mount layer-specific transcriptional responses to mechanical stress.

We previously showed that early seed elongation is driven by a mechanosensitive response in the ab-oi layer that aligns cortical microtubules (CMTs) along shape-induced stress patterns^17^. We therefore tested whether the early elongation defects of *ap2* seeds reflected altered CMT organization. At 2DAP, CMTs were highly ordered and aligned with the seed’s main curvature axis, in both WT and *ap2-6* mutant (**Figure 6D-E**). By 5DAP, CMTs were still preferentially oriented according to the main curvature axis in both genotypes (**Figure S6D**), but they remained highly organized in *ap2-6*, whereas they became disorganized in WT (**Figure 6F**). These data indicate that *AP2* activity is not required for the mechanosensitive alignment of the CMTs according to shape-driven stresses in the ab-oi layer, but is needed for them to become disorganize over time, a transition that normally accompanies the shift from anisotropic to isotropic growth in WT seeds^17^.

Because *ap2* seeds cannot be grown *in vitro* for long time periods, we could not directly assess how CMT organization affects growth anisotropy in the mutant. Instead, we compared ab-oi cell morphology in seeds at different developmental stages. In WT seeds, ab-oi cells gradually became isotropic in shape over time, consistent with the CMT disorganization observed in this layer. In contrast, ab-oi cells in *ap2-6* remained anisotropic in shape, forming concentric rings of elongated cells oriented perpendicularly to the seed radius (**Figure 6G**), in agreement with the CMT pattern observed in the ab-oi layer. This pattern differed markedly from that of the *katanin1* (*ktn1*) mutant, in which reduced CMT responsiveness also leads to reduced seed elongation (**Figure S6E**). These comparisons suggest that the round shape of *ap2* seeds does not arise from impaired CMT mechanosensitivity but from a shift in the direction of cell growth in the ab-oi layer after fertilization, being axial in the WT but radial in the *ap2-6* mutant.

Altogether, these results demonstrate that AP2’s primary role in seed growth lies in the control of wall composition after fertilization as an inherent component of specific differentiation programs, rather than through regulating tissue-specific mechanosensitive responses.

## Discussion

Seed size and shape arise from the coordinated interplay of developmental and mechanical processes within the seed coat^14,16,17^. Here, we identify APETALA2 (AP2) as a key regulator that links outer integument differentiation with the control of cell wall composition and growth mechanics after fertilization.

A central finding of our study is that AP2 regulates pectin and hemicellulose patterning within the outer integument walls. Loss of *AP2* function disrupts the spatial distribution of pectin methylesterification in each outer integument layer and reduces xyloglucan accumulation, leading to altered wall thickness and compromised cell integrity. Surprisingly, AP2 activity also strongly affects pectin acetylation, which increases throughout development in WT seeds but not in *ap2-6* mutants. Although the spatial pattern of this modification could not be probed due to a lack of antibody, this type of pectin modification, which is known to affect the properties of MSC walls^54^, may, together with the demethylesterification, promote changes in pectin properties that are also relevant for seed growth control^55,56^. Nevertheless, genetic perturbations affecting pectin methylesterification and hemicellulose biosynthesis reproduce specific aspects of the *ap2* growth phenotype, supporting a direct connection between cell wall composition and the dynamics of seed expansion. These results are consistent with recent studies reporting seed growth defects in other pectin- and hemicellulose-associated mutants^57–59^. Thus, AP2 fine-tunes cell wall material properties to balance expansion and mechanical constraints during seed development. This function may not only be essential for proper seed development but also for adjusting seed size in response to environmental signals, as AP2 was recently shown to be repressed by *CONSTANS*, thereby allowing seed growth to adapt to photoperiod conditions^60^.

Despite its strong influence on wall architecture, *AP2* appears dispensable for mechanosensitive transcriptional and cytoskeletal responses. Both *ELA1* and *H1.3* reporters remained inducible by compression in *ap2* mutants, indicating that mechano-perception and -transduction into tissue-specific responses persist in the absence of AP2 function. Moreover, cortical microtubules (CMTs) were properly aligned with predicted stress orientations in *ap2-6*, suggesting that the seed shape defects of this mutant do not stem from reduced CMT mechanosensitivity. One possible explanation is that these defects arise from the altered shape of *ap2-6* ovules, which may impose an initial stress pattern that orients CMTs differently compared with the WT. However, seed elongation is also observed in the *ap2-5* mutant, whose ovules resemble those of other *ap2* alleles. An alternative hypothesis is that seed elongation is driven by specific changes in wall properties that occur after fertilization in WT seeds but not in *ap2* mutants, and that these differences are subsequently amplified through CMT responses to mechanical forces, similar to the situation described in the hypocotyl, where asymmetric pectin demethylesterification of epidermal walls breaks symmetry to initiate organ elongation^61^.

Beyond its implications for the mechanical control of organ growth, our work also provides new insights into the molecular regulation of seed coat identity. It indeed supports a two-step model in which each integument layer acquires an initial identity during ovule development that is subsequently refined after fertilization, a process that requires, at least in part, *AP2* activity. This refinement could depend on auxin, as this hormone is transported from the endosperm to the seed coat to promote its development after fertilization by alleviating PRC2-dependent repression of gene expression and enabling the recruitment of JUMONJI-type (JMJ) histone demethylases to specific loci^62,63^. How these mechanisms contribute to the differentiation of each integument layer into distinct cell types, and how they intersect with AP2 activity in the outer integument, remain to be determined. Investigating this interplay could yield important insights into how a key regulator of cell identity and differentiation exerts distinct roles throughout tissue development.

Finally, given the marked diversity in marker gene expression patterns observed in WT and *ap2-6* ovules and developing seeds after fertilization, the outer integument emerges as a powerful model for studying cell identity establishment and maintenance throughout organogenesis. The mechanisms controlling epidermis-associated gene expression in the seed coat are of particular interest. Epidermal identity markers, which are normally restricted to the outermost cell layer of organs^12,34,35^, are expressed in all integument layers during ovule development, suggesting that lineage information overrides positional cues at this stage. However, this situation changes after fertilization, when lineage cues appear to be overridden so that only the abaxial outer integument (ab-oi), and to a lesser extent the adaxial outer integument (ad-oi), maintain expression of epidermis-associated genes and differentiate using molecular pathways, such as the MBW pathway, that are usually involved in the differentiation of epidermal sub-populations^40,64^. Although *AP2* activity provides positional information necessary for *GL2* and *PDF1* expression in the ab-oi layer, likely through the MBW pathway, it does not account for the post-fertilization expression patterns of *ATML1* and *PDF2*, implying that additional factors contribute to outer integument identity in addition to AP2. Expression of epidermal identity genes at organ surfaces was recently shown to depend on the transduction of mechanical cues^10,65^, which remain active in *ap2*. Integrating novel findings relating to mechanical signaling outputs with AP2-dependent regulation of outer integument identity could thus open new avenues for understanding how biochemical and mechanical signals interact to coordinate cell identity and organ morphogenesis.

## Materials and Methods

### Plant Material and Growth Conditions

Information on the mutants and transgenic lines used in this study, all in the *Arabidopsis thaliana* Columbia-0 (Col-0) background, is provided in **Table S15**.

For experiments involving only wild-type (WT) plants, seeds were surface-sterilized in 70% ethanol containing 0.05% Triton X-100 (Sigma) for 15 min, rinsed three times in 95% ethanol, and dried on Whatman paper for 30–60 min under a sterile hood. Sterilized seeds were sown on plates containing 1× Murashige and Skoog (MS) medium (pH 5.7, with antibodies for selection of the transgenic lines) and stratified for 2 days at 4°C in the dark. Plates were then transferred to a growth chamber (Sanyo) under short-day conditions (8 h light; 21°C day / 18°C night). For experiments involving WT and *ap2* mutant plants, which are sensitive to sterilization, seeds were sown directly on soil (Argile 10, Favorit), stratified for 2 days at 4°C in the dark, and germinated under short-day conditions (8 h light; 21°C day / 19°C night) inside plastic bags. After two weeks, seedlings (from both soil and MS medium) were transferred to individual pots containing soil and grown for one week under short-day conditions before being moved to long-day conditions (16 h light; 21°C day / 19°C night) for the remainder of their life cycle.

Around one week after bolting, flowers from the main stem were manually pollinated at anthesis. All plants were self-pollinated except *ap2* mutant alleles, that were always pollinated with WT pollen to maintain homozygosity in maternal tissues only.

### Generation of Transgenic Lines

#### *pATML1::mCitrine-ATML1* in *atml1-3* mutants

The *pATML1::mCitrine-ATML1* vector was constructed by VectorBuilder with the vector ID VB210104-1143ajf. The vector consists of 6145 bp upstream of the *ATML1* gene that includes the promoter region, followed by the mCitrine fluorophore sequence attached by a linker to the *ATML1* gene including introns. 985 bp downstream of the ATML1 gene was included to provide the ATML1 terminator. The binary vector contains the Hygromycin B plant resistance cassette. The bacterial resistance is Kanamycin. *atml1-3* homozygous plants were transformed with the *pATML1::mCitrine-ATML1* vector by the agrobacteria-mediated floral dip method^66^. T1 plants were selected for Hygromycin resistance. A line was selected that segregated 3:1 as a single locus and which rescued the number of sepal giant cells, suggesting it was expressed at nearly endogenous levels.

#### *pPDF2::mScarlet-PDF2 in pdf2-4* mutants

The *pPDF2::mScarlet-PDF2* plasmid was constructed using NEBuilder HiFi DNA Assembly following the protocols of *New England BioLabs*. 6,147 bps upstream of the *PDF2* gene was PCR amplified from genomic DNA with overhangs for ligation. Additionally, the *PDF2* gene including exons and 1,551 bps downstream of *PDF2* was PCR amplified from genomic DNA with overhangs for assembly. The mScarlet gene was PCR amplified from the mScarlet-mTurquoise plasmid (Addgene 98839) with an AAA Kozac consensus sequence at the 5’ end and linker sequences at the 3’ end as well as overhangs for assembly. These DNA fragments were assembled with NEBuilder into the vector pART27, which confers Spectinomycin resistance in bacteria and Kanamycin resistance in plants. The assembly reaction was electroporated into competent SMC4 *E.coli* cells. The plasmid was sequence verified and mated into *Agrobacterium tumefaciens* strain GV3101. *pdf2-4* homozygous plants were transformed with the *pPDF2::mScarlet-PDF2* vector by the agrobacteria-mediated floral dip method^66^. T1 transformants were selected on Kanamycin. A line was selected in which segregated 3:1 as a single locus and which rescued the number of sepal giant cells, suggesting it is expressed at nearly endogenous levels.

#### *pGL2::3X-VENUS-N7* in Col-0

The *GL2* promoter (1,571 bp upstream of the coding sequence) was amplified by PCR (primers in **Table S16**) and assembled into a shuttle vector containing the *3X-VENUS-N7* reporter, linearized with KpnI and BamHI, using NEBuilder HiFi DNA Assembly. Assembly products were transformed into *E. coli* TOP10 cells, selected on ampicillin (100 µg/mL), and screened by PCR and sequencing. The shuttle and *pML-BART* binary vectors were both digested with *NotI* and ligated using T4 DNA ligase (NEB). Positive *E. coli* clones were selected on spectinomycin (100 µg/mL) and X-GAL, verified by PCR, and used for *Agrobacterium*-mediated transformation. Col-0 plants were transformed by floral dipping^66^, and T₁ plants were selected on BASTA. Several single-locus lines (3:1 segregation in T₂) were advanced to homozygosity (T₃). T₂ plants from two independent lines were crossed with *ap2-6* to generate *ap2-6 × pGL2::3X-VENUS-N7*.

#### *pAP3::3X-VENUS-N7* and *pGILT::3X-VENUS-N7*

Promoter regions of *AP3* (998 bp upstream) and *GILT* (3,000 bp upstream) were amplified (primers in **Table S12**) and cloned into the *pML-BART* binary vector using the same two-step cloning procedure as described for *pGL2::3X-VENUS-N7*. Col-0 plants were transformed by floral dipping^66^ and selected for BASTA resistance. Single-locus lines (3:1 segregation in T₂) were advanced to homozygosity (T₃).

### Morphometric Analysis

For most experiments, siliques were opened with a needle at anthesis or specific time points after pollination. Seeds were placed in clearing solution (1 vol glycerol / 7 vol chloral hydrate; *VWR*) between a slide and coverslip and incubated for ≥24 h at 4°C before imaging on a Zeiss Axioimager 2 microscope with a 20× DIC objective. For the second *PMEI-OX* replicate, siliques were fixed in ethanol/acetic acid (9:1, v/v) for 24 h at 4°C, washed in 90% ethanol for 10 min, and stored in 70% ethanol before chloral hydrate clearing with and imaging. Seed size and shape were semi-automatically quantified on ImageJ using a dedicated script as described previously^17^.

### RNA Sequencing and Analysis

Total RNA was extracted from developing seeds at 2 and 5 days post-anthesis (DPA) (10 replums per replicate; 3 biological replicates) using the Spectrum Plant Total RNA Kit (Sigma-Aldrich) and treated with Turbo DNA-free DNase I (Invitrogen). RNA quality was verified using a Dropsense spectrophotometer (PerkinElmer).

Library preparation, sequencing, and analysis were performed at the IPS2 Transcriptomics Platform (POPS). RNA integrity and quantity were assessed using an Agilent Bioanalyzer and the RiboGreen kit (Thermo Fisher). Libraries were prepared using Illumina protocols and sequenced (2 × 150 bp) on a NovaSeq (Illumina). Reads were mapped to the *A. thaliana* TAIR10 genome (release 42), and differential expression analysis was performed using DicoExpress^67^. Genes with low counts were filtered using the “filterByExpr” function where the group argument specifies the biological conditions, the min.count value is set to 15 and the other arguments are set to their default values. Libraries were normalized with the TMM method with the default parameter values. The differential analysis was based on a negative binomial generalized linear model in which the logarithm of the average gene expression is an additive function of a genotype effect (*ap2-6*, Col0), a DAP effect (2, 5), their interaction and a replicate effect (3 modalities). We tested the significance of the difference between *ap2-6* and Col0 at 2 and 5 DAP using a likelihood ratio test. Raw p-values were adjusted with the Benjamini–Hochberg procedure to control the false discovery rate.

Except specified otherwise, differentially expressed genes (DEGs) were defined as |log₂FoldChange| ≥ 2 and an adjusted p-value ≤ 0.05. GO term enrichment analysis and representation were done using the version v0.741 of [ShinyGO](https://bioinformatics.sdstate.edu/go74/). Seed-compartment expression patterns were examined using BAR’s [eNorthern tool](https://bar.utoronto.ca/). Reclustering of the data and heatmap representation according to Euclidean distance were performed in R. Measurements of the overlaps between *ap2-6* DEGs at 2 and 5 DAP and the genes enriched in ab-oi or ad-oi layers at 3 and 5 DAP (data from^30^) were performed using [Venny](https://bioinfogp.cnb.csic.es/tools/venny/) and statistical significance was calculated using Fisher’s Exact Test for Count Data on R, assuming 27,880 *Arabidopsis* genes found in at least one sample of our transcriptomic dataset.

### qPCR Analysis

RNA extraction and DNase treatment were performed as described above. cDNAs were synthesized using the SuperScript VILO cDNA Synthesis Kit (Invitrogen) and diluted 1:20 in Milli-Q water before qPCR. Reactions were run in 384-well plates on a QuantStudio™ 6 Flex system (Thermo Fisher) using FastStart Universal SYBR Green Master (ROX) (Roche) in 10 µL total volume. Cycling conditions were 95°C for 10 min, followed by 40 cycles of 95°C for 10 s and 60°C for 30 s. Data were analyzed using QuantStudio Real-Time PCR Software v1.3. Gene expression was normalized to the geometric mean of *EIF4A1* and *AP2M*. PCR efficiency (E) was calculated from standard curves (E = 10^(-1/slope)), and expression levels were reported as E^(–ΔCt), where ΔCt = Ct_GOI – Ct_REF. Primer sequences are listed in **Table S16**. Each experiment included five biological replicates per genotype and condition, and was repeated twice with independent plant batches.

### Histology

Histology was performed as described previously^16^. Seeds were fixed in ice-cold PEM buffer (50mM PIPES, 5mM EGTA and 5mM MgSO4, pH 6.9) with 4% (w/v) paraformaldehyde. The samples were placed under vacuum (2 x 30 min on ice), rinsed twice in PEM buffer, dehydrated through an ethanol series and infiltrated with increasing concentrations of LR White resin in absolute ethanol (London Resin Company) over 8 days before being polymerized at 60°C for 24h. The samples were sectioned (1.0 µm thickness) using a diamond knife 45° angle (Diatome, LFG Distribution) mounted on a Leica RM6626 microtome and dried onto glass slides.

For toluidine blue staining, the sections were incubated for 20 seconds at 70°C with filtered Toluidine Blue 1% / 1% borax before being rinsed with distilled water, dried and mounted in Entellan mounting medium (Merck). The sections were imaged with a Zeiss Axioimager 2 equipped with a 20x dry objective. Images were analyzed using the ImageJ software.

For immunostaining with LM19, JIM5 and LM15, the sections were blocked with 3% w/v BSA in 1X PBS and incubated for 1 h at room temperature. For LM20, an incubation in CAPS buffer (50 mM CAPS and 2 mM CaCl_2_) for 1 h followed by three washes with 1x PBS was performed before blocking with 3% skimmed milk in 1X PBS. For LM20, an incubation in CAPS buffer (50 mM CAPS and 2 mM CaCl2) for 1 h followed by three washes with 1x PBS was performed before blocking with 3% skimmed milk in 1X PBS. For LM25 staining after pectolyase treatment, an incubation with 0.1% pectolyase in 1X PBS treatment was done for 25 min at room temperature and washed once in 1X PBS before the blocking step. The antibodies were applied to the sections overnight at 4°C in a humid chamber. The JIM5, LM15 and LM19 antibodies were diluted 1:10 (v/v) in PBS/BSA 1%. The 2F4 antibody was diluted 1:5 in 1% skimmed milk in TCaS buffer. The LM20 antibody was diluted 1:10 (v.v) in CAPS buffer (50 mM CAPS and 2 mM CaCl_2_). The sections were then washed in an excess of the buffer to dilute the antibody and subsequently incubated for 1 hour at room temperature with the secondary antibody (anti-rat IgG Alexa 488, anti-rat IgM Dylight Alexa 488, and anti-mouse IgG Alexa 488 for JIM5/LM15, LM19/LM20, and 2F4, respectively) diluted 1:100 in the same buffers as those used for diluting the primary antibody. The sections were washed in buffer solutions as described above and covered with their respective buffer. The samples were then counterstained with filtered Calcofluor White M2R (fluorescent brightener 28; Sigma-Aldrich) at 0.25 mg.ml^-1^ and mounted with VECTASHIELD (Eurobio). The sections were imaged using a Zeiss Axioimager 2 equipped with a 40x dry objective. Images were analyzed using the ImageJ software. Intensity color coding was done using the Fire lookup table.

### Confocal Microscopy and Image Analysis

Mature carpels at anthesis and developing siliques at specific stages of seed development were opened with a needle, and seeds were mounted on double-sided tape in water or ½ MS solution. In some cases, cell walls were stained with propidium iodide (Sigma, 1:10 dilution, 10 min) or CarboTag-AF488 (Joris Sparkel^68^, 100 µM, 30 min) and washed twice in ½ MS before imaging. Fluorescent reporters were imaged using Leica SP8 or Zeiss LSM980 confocal microscopes with 25× water-dipping objectives. Z-stacks (512×512 or 1024×1024 pixels) were collected with 1–2 µm z-steps.

Z-stacks were analyzed with ImageJ. Maximal projections and z-section (5 to 10 slices thick) were performed, and intensity color coding was done using the Fire lookup table. Microtubule imaging was done on z-stacks from the *p35S::MAP65-1-*RFP reporter obtained with a Zeiss LSM980 confocal microscope using the Airyscan mode for high resolution imaging, as described in^17^. The analysis of microtubule organization and correlation with seed curvature was performed using the FibrilTool plugin^69^ on [MorphographX](https://morphographx.org/) according to^17^. Analysis of ab-oi cell size and aspect ratio was performed on WT and mutant seeds expressing the *LTi6b-GFP* or *MAP65.1-RFP* reporters and using the [MorphographX](https://morphographx.org/) software according to^17^. The analysis of the integrity defects of outer integument cells of seeds at 5DAP was performed manually on WT and mutant seeds expressing the *LTi6b-GFP* or *MAP65.1-RFP* reporters with ImageJ.

For the quantitative analysis of the *pH1.3::H1.3-GFP* and of the *pELA1::3X-VENUS-N7* reporters, nuclei were automatically detected in seed confocal images as local maxima in a Gaussian scale-space, using standard deviations varying from 1.0µm to 2.0µm. The fluorescent signal intensities were then quantified for each detected nucleus as a Gaussian-weighted average of the voxel intensities around the detected points (Sigma = 1µm) to obtain a value per nucleus of the considered seed. These analyses were carried out through a [Python script](https://gitlab.inria.fr/gcerutti/seednucleifluo) built using the [Gnomon](https://gnomon.gitlabpages.inria.fr/gnomon) computational platform and relying on plugins from the [Tissue Image ToolKit library](https://mosaic.gitlabpages.inria.fr/timagetk/).

For the analysis of ab-oi cell death in seeds grown *in vitro*, isolated WT and *ap2-6* seeds expressing *LTi6b–GFP* (replicate 1) or *LTi6b–tdTomato* (replicate 2) were placed on double-sided tape, and covered with a ½ MS solution containing 1% sucrose (Sigma), 1× Gamborg Vitamins (Sigma), and 0.1× PPM (Plant Preservative Medium, Plant Cell Technology), supplemented with different concentrations of sorbitol (0, 200, or 500 mM, Sigma). Each seed was imaged at 0 h to assess its integrity at the start of the experiment and again at 5 h to evaluate survival during cultivation. Cell death in the ab-oi layer was assessed manually using ImageJ.

### Transmission Electron Microscopy

Seeds were fixed in 4% formaldehyde and 2% glutaraldehyde in 0.1 M phosphate buffer (PB) (pH 7.2) under vacuum (0.6 bar) for 1 h at 4°C, then incubated overnight in fresh fixative. Samples were washed 3 times in PBS, post-fixed in 1% osmium tetroxide in 0.1M PB (pH 7.2) for 2 h at RT, rinsed 3 times in 0.1M PB (pH 7.2) and 3 times in H20. After dehydration through a graded ethanol series (25 to 96% vol / vol) with 20-min incubations under vacuum in each bath at RT and 2 times in 100% ethanol during 1 h under vacuum, the samples were infiltrated with low-viscosity Spurr resin (Electron Microscopy Sciences) using a graded series in ethanol (33, 66, and 2 x 100% vol/vol Spurr resin) at RT, with each incubation lasting 24 h (including 1 h under vacuum). The samples were polymerized in fresh Spurr resin at 60 °C for 18 h. Ultrathin sections (70 nm) were prepared using UC7 Leica Ultramicrotome, placed on formvar-coated grids. Sections were examined under JEOL 1400 TEM at 120 kV and imaged with a Gatan Rio 16 camera. Images were analyzed in ImageJ, and cell wall thickness was measured manually at three positions per cell wall in at least 3 seeds per genotype.

### Application of Mechanical Constraints

Mechanical compression was applied to 1 DPA siliques using a microvice for 24 h under growth chamber conditions, as described previously^15^. Compressed seeds were processed and imaged as described above. Optical sections of 3D stacks were generated on ImageJ to confirm efficient compression.

### Cell Wall Enzymatic Fingerprinting

Cell wall fingerprinting was performed as described previously^70^. Seeds from 10 siliques per replicate (three biological replicates per stage and genotype) were dissected, immersed in 96% ethanol, and dried overnight at 30°C in a SpeedVac. Dry seeds were digested with 5 U/mg dry weight of endo-polygalacturonase (Megazyme) or of xyloglucanase (Megazyme) in 50 mM ammonium acetate buffer (pH 5) at 37°C for 18 h. Supernatants were collected after centrifugation (13,000 rpm, 10 min), and 10 µL was injected for LC–MS analysis using an UltiMate 3000 RS HPLC system (Thermo Fisher Scientific) coupled to an Impact II UHR-QqTOF mass spectrometer (Bruker Daltonics) equipped with an electrospray ionization (ESI) source in negative mode (capillary voltage 4,000 V; nebulizer 40 psi; dry gas 8 L/min; 180°C). Chromatographic separation was performed on an ACQUITY UPLC Protein BEH SEC column (125 Å, 1.7 µm, 4.6 × 300 mm; Waters) with a guard column, using 50 mM ammonium formate and 0.1% formic acid at 0.4 mL/min and 40°C. Data were processed in Compass 1.8 (Bruker Daltonics) as described previously^71^.

### Statistical analysis

Unless stated otherwise, all experiments were conducted at least two times on two independent batches of plants. Data from these experiments were either pooled or presented separately in distinct graphs in main and in supplementary figures, as appropriate. For each genotype or condition, seeds were collected from at least three fruits originating from three different plants. The number of seeds (biological replicates) analyzed in each experiment was determined based on technical constraints such as time and material availability. No sample size calculations were performed prior to the experiments. Seeds that were severely damaged during sample preparation (particularly *ap2* seeds) were excluded from analysis. Data analysis was performed using Microsoft Excel (v.2016), R (v.4.3.2), or Python (v.3.7.5). For comparisons between two groups, bilateral Student’s tests were used. For comparisons between multiple groups, one-way ANOVA with Tukey’s multiple comparison tests were used. For RT-qPCR experiments, the groups were compared using a Kruskal-Wallis test followed by Dunn’s Post-hoc test. Frequency comparisons were done with Chi² tests. Statistical significance was indicated with asterisks according to the following convention: p < 0.05 (*), p < 0.01 (**), p < 0.001 (***), and p < 0.0001 (****).

## Data availability

All the raw images and data shown in this study will be deposited on [Zenodo](https://zenodo.org/) upon publication. The transcriptomic dataset has been deposited on the [Gene Expression Omnibus platform](https://www.ncbi.nlm.nih.gov/geo/) and will be made public upon publication

## Acknowledgements

We kindly thank George Coupland for providing the *pAP2::AP2-VENUS* reporter, Yvon Jaillais for the *LTi6b-GFP* reporter, Charlotte Kirchhelle for the *LTi6b-tdTomato x RABa5c-YFP* reporter, Olivier Hamant for the *MAP65-1-RFP* reporter, Françoise Monéger for the *PMEI4* and *PMEI5* overexpressors, Jan Traas for the *xxt1 xxt2* double mutant, and Stéphane Verger for the *ktn1* mutant. We warmly thank Joris Sprakel and his group for providing the CarboTag-AF488 dye. We acknowledge the contribution of SFR Biosciences (Universite Claude Bernard Lyon 1, CNRS UAR3444, Inserm US8, ENS de Lyon) PLATIM-LyMIC, of the Bio21 Advanced Microscopy Facility (University of Melbourne), and of Claire Lionnet from RDP for technical assistance with microscopy; of Alexis Lacroix, Patrice Bolland, Camille Knaupp, Joseph Pacitto and Justin Berger for technical assistance with plant cultivation; of Isabelle Desbouchages and Hervé Leyral for technical assistance regarding molecular biology work; of Cindy Vial, Laureen Grangier, Nelly Camilleri, Stéphanie Maurin and Julie Prata for administrative assistance; of Jeremy Just, Nicolas Doll, and Marie-Laure Martin for help with transcriptomic analysis, and of the IJPB’s Plant Observatory Platforms (PO-Chem) for the enzymatic fingerprinting. The PhD thesis of Camille Bied was supported by a fellowship from the French Ministry of Higher Education. The PhD thesis of Amélie Bauer and of Runjue Yao were supported by a joint PhD program between the CNRS and the University of Melbourne. This work was also supported by the French National Agency of Research (ANR, grant agreement ANR-23-CE13-0009, “MECHASEED”, ANR-22-CE43-0013, “WALLDERIVE”), by the research fund of the *ENS de Lyon*, by the *BAP* department at *INRAE*, and by the European Research Council (ERC, grant agreement No 101019515, “MUSIX”).

## Author contributions

Design of the work: CB, AV, GI, JG, BL

Acquisition of data: CB, RY, AC, LL, AB, SB, JB, ALA, AV, BL

Analysis of data: CB, RY, AC, LL, AB, JB, CPLR, AV, BL

Creation of new genetic material used in this work: CB, RY, DD, FC, AR, BL

Creation of new software used in the work: GC

Drafting of the work: CB and BL, with inputs from all co-authors

---

**Figure S1.**
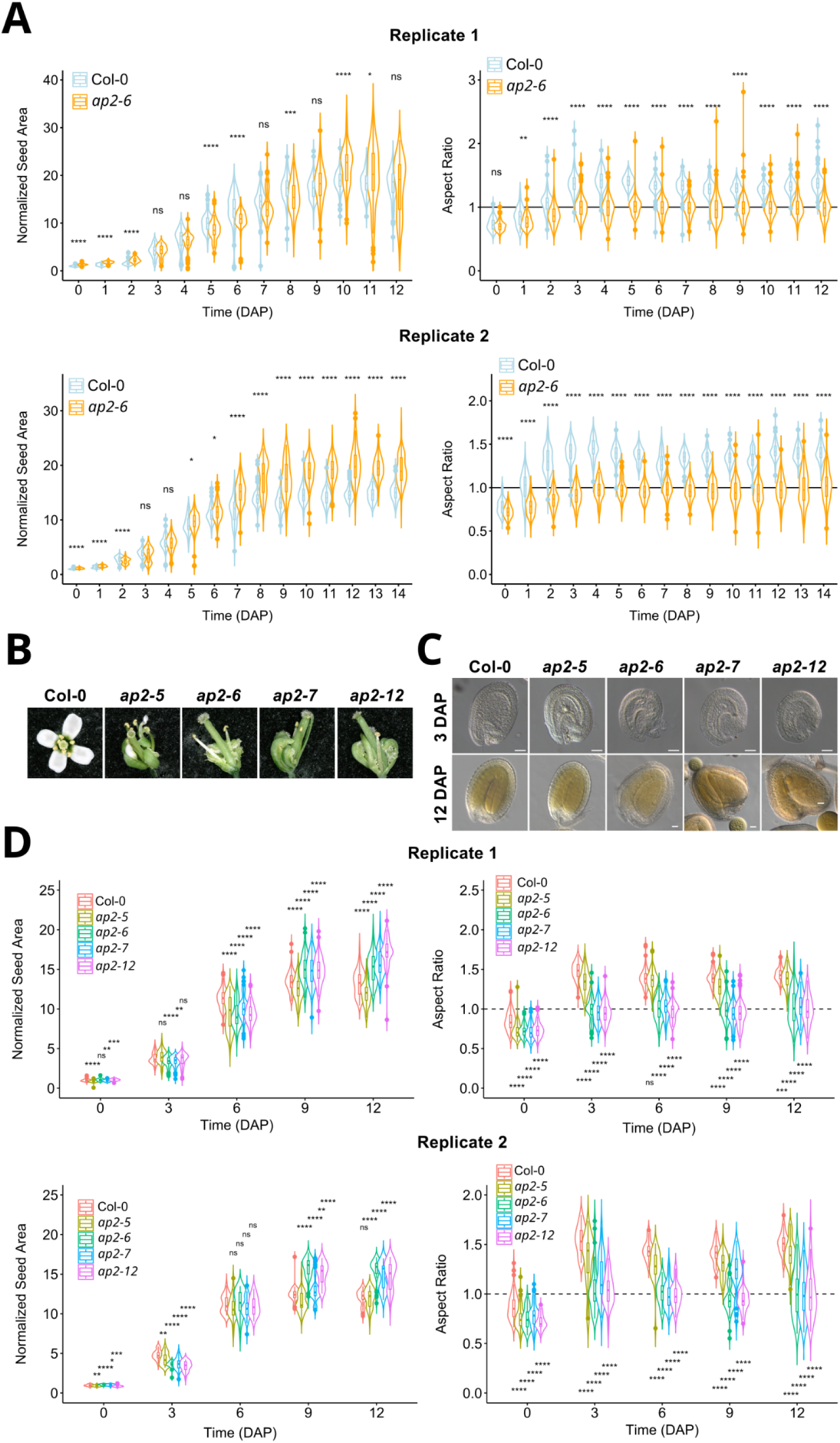
Analysis of the growth dynamics of *ap2* alleles of increasing strengths. **A.** Growth dynamics of WT and *ap2-6* seeds (crossed with WT pollen) from 0 to 12 - 14 days after pollination (DAP) from two experiment independent from the shown in Figure 1; replicate 2: n = 34 - 123 seeds from 3 fruits per day per genotype; replicate 3: n = 65 – 139 seeds from 3 fruits per day per genotype. Data were compared using bilateral Student’s tests. **B** Representative WT and *ap2* mutant flowers showing homeotic conversions of the floral organs in *ap2* mutant alleles. **C.** Representative WT and *ap2* mutant seeds (crossed with WT pollen) at 3 and 12 DAP, scale bars, 100 µm. **D.** Growth dynamics of WT and *ap2* mutant seeds (crossed with WT pollen) seeds from 0 to 12 DAP; two independent experiments; replicate 1: n = 71 – 123 seeds from 3 fruits per day per genotype; replicate 2: n = 20 – 124 seeds from 3 fruits per day per genotype. Data were compared using bilateral Student’s tests.

**Figure S2.**
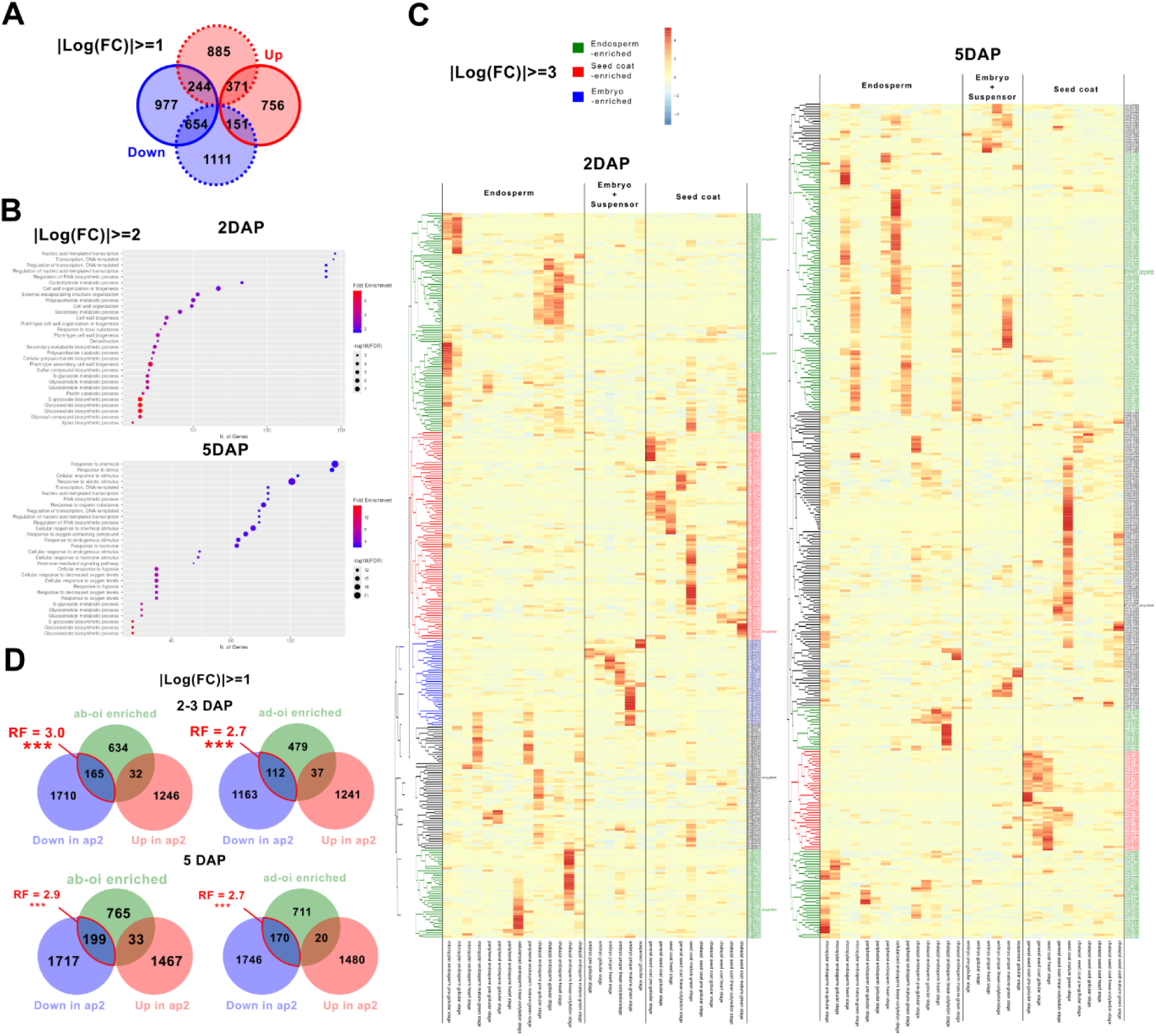
Analysis of *ap2-6* Differentially Expressed Genes (DEGs) **A**. Number of DEGs down or up-regulated at 2 and 5 DAP in *ap2-6* seeds (crossed with WT pollen) compared to the WT (|log₂FC| ≥ 1, *p* ≤ 0.05). **B.** Graphic representation of the 30 most enriched Gene Ontologies (GOs, calculated according to their p-value) in *ap2* DEGs (|log₂FC| ≥ 2, *p* ≤ 0.05) at 2 and 5 DAP, obtained using ShinyGO. **C.** Heatmap showing the clustering of *ap2-6* strongest DEGs at 2 and 5 DAP based on their expression in the seed compartments according to the dataset from^2^. Genes highlighted in green, blue and red, are part of gene clusters whose expression is enriched in endosperm, embryo, and seed coat, respectively. Note that a higher cutoff (|log₂FC| ≥ 3, *p* ≤ 0.05) was used in these heatmaps to reduce gene number and simplify visualization. **D.** Overlap between the genes enriched in the ab-oi or ad-oi layer at 3 and 5 DAP (|log₂FC| ≥ 1, *p* ≤ 0.05, data from^1^) and the genes under and over-expressed in *ap2-6* at 2 and 5 DAP (|log₂FC| ≥ 1, *p* ≤ 0.05), respectively. Each representation factor (RF) and its significance was calculated using a Fisher’s Exact Test for Count Data.

**Figure S3.**
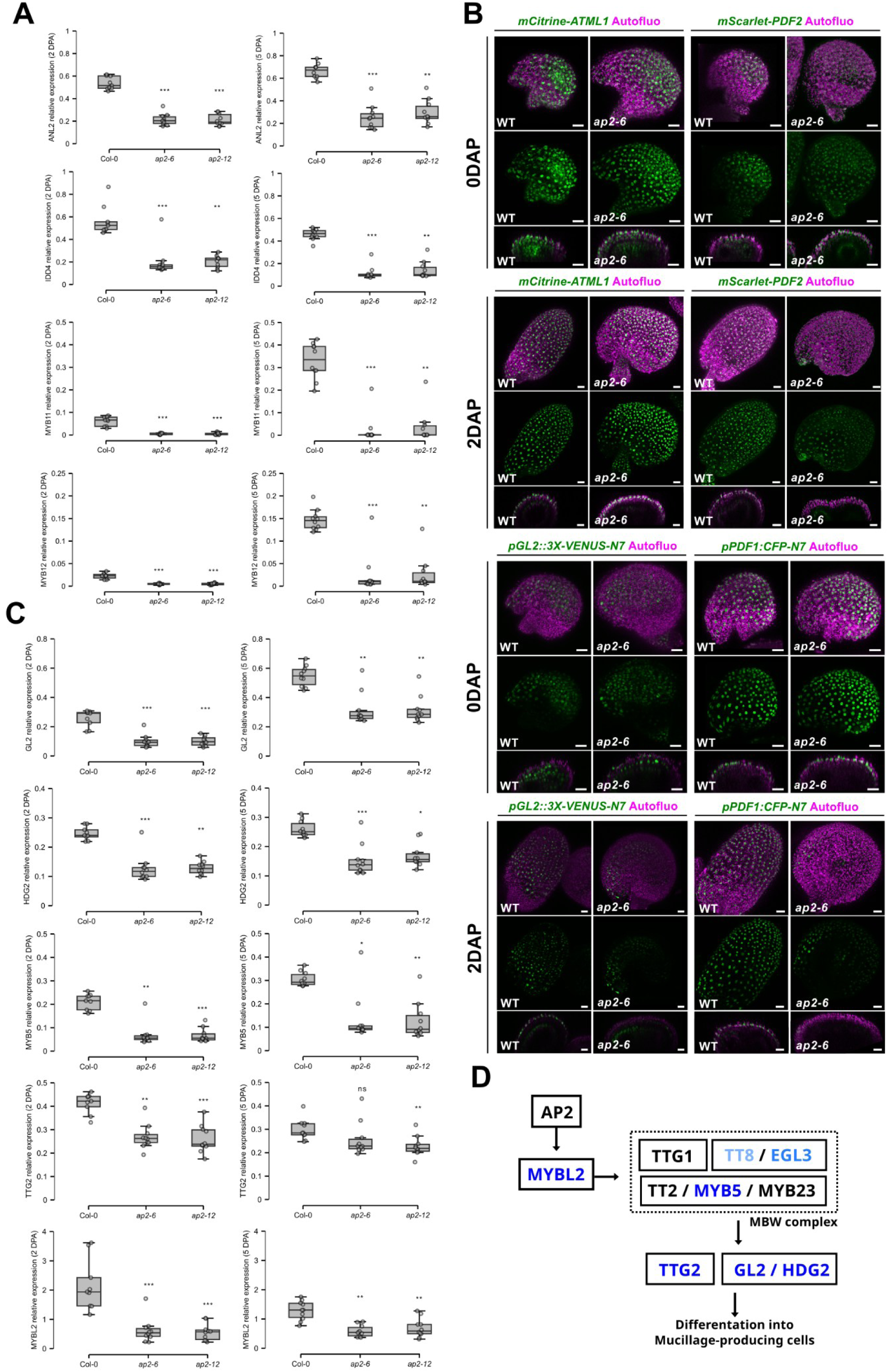
Analysis of the Gene Regulatory Network acting downstream of *AP2* in the control of outer integument differentiation. **A**. Relative expression of *ANL2, IDD4*, *MYB11* and *MYB12* at 2 and 5 DAP in the WT and in the *ap2-6* and *ap2-12* mutants (crossed with WT pollen); n = 10 replicates from 2 independent experiments. Statistical differences established using the Kruskal-Wallis test followed by Dunn’s Post-hoc test. Expression of these TFs is enriched in the ad-oi layer according to^1^ and *ANL2, IDD4*, *MYB12* are predicted to be direct targets of AP2 according to^3^. **B.** Representative projections and z-slices of mature ovules (0 DAP) and developing seeds (2 DAP) of WT and *ap2-6* seeds (crossed with WT pollen) expressing transcriptional reporters for *GL2* (*pGL2::3X-VENUS-N7*) and *PDF1* (*pPDF1::CFP-N7*), and translational reporters for *ATML1* (*pATML1::mCitrine-ATML1*) and for *PDF2* (*pPDF2::mScarlet-PDF2*). *GL2:* n = n= 30 – 45 seeds from 6 - 7 plants, two independent experiments; *PDF1*: 35 – 37 seeds from 7 – 8 plants, two independent experiments; *ATML1*: n = 15 seeds from 3 plants, one experiment; *PDF2*: n= 14 – 19 seeds from 3 – 4 plants, one experiment; Scale bars, 10 µm. **C.** Relative expression of members of the *MYB-BHLH-WD40* (*MBW*) pathway (*GL2*, *HDG2*, *MYB5* and *MYBL2*) in the WT and in the *ap2-6* and *ap2-12* mutants (crossed with WT pollen); n = 10 replicates from 2 independent experiments. Statistical differences established using the Kruskal-Wallis test followed by Dunn’s Post-hoc test. **D.** Model of *AP2* regulation of ab-oi differentiation into mucilage-producing cells according to^4–6^. Genes labelled in blue are downregulated in *ap2* in our datasets.

**Figure S4.**
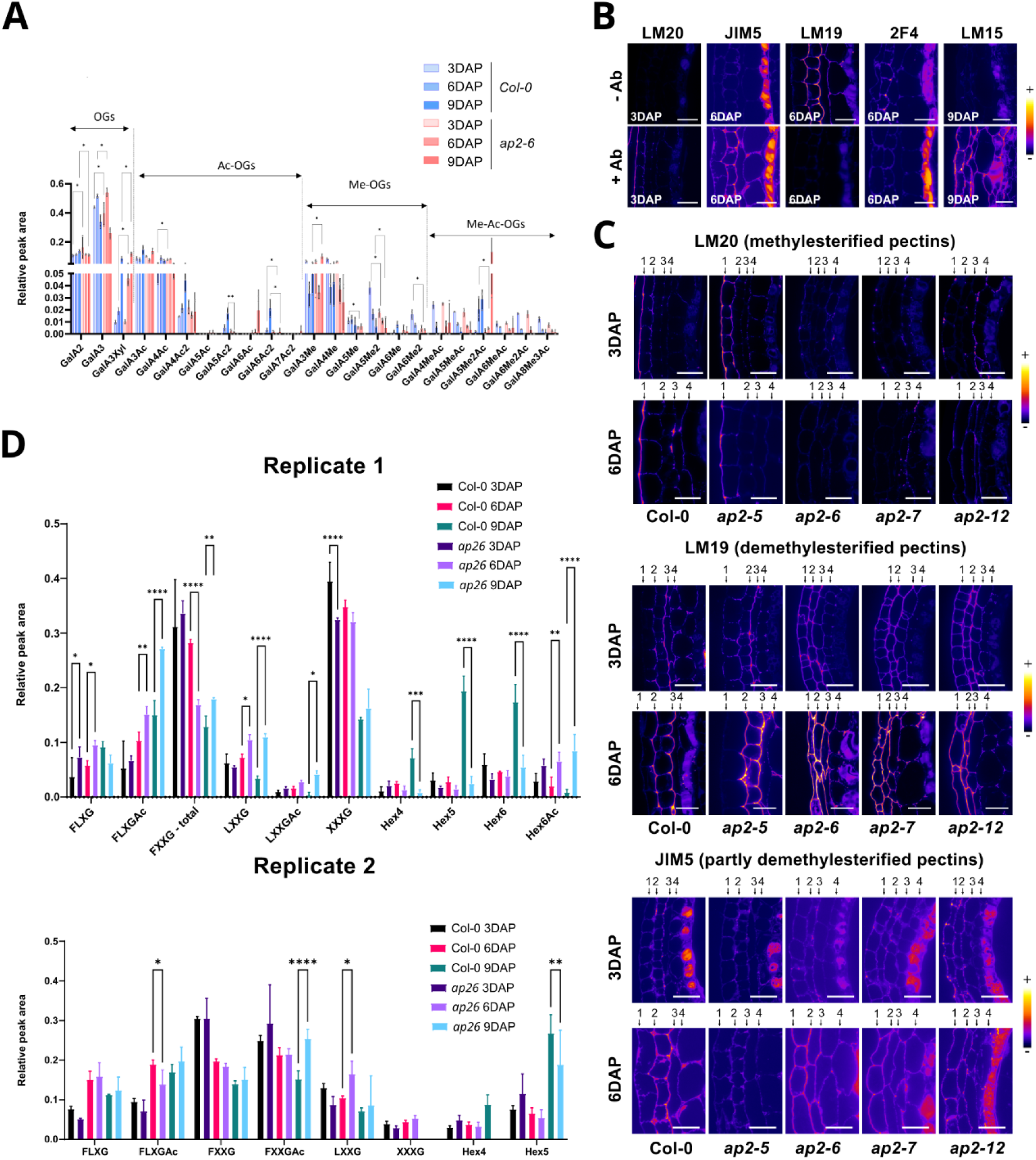
AP2-dependent regulation of seed coat wall composition. **A.** Relative level of the different oligogalacturonides released following digestion of intact seeds at 3, 6 and 9 DPA with a polygalacturonase. 1 experiment with 3 technical repeats independent from the one presented in Fig.4, WT and *ap2-6* (crossed with WT pollen) were compared using bilateral Student’s tests. **B.** Verification of the specificity of the staining of WT seeds with the LM20, LM19, JIM5, 2F4 and LM15 (with pectolyase treatment) antibodies through a comparison between a section treated with the antibody (+Ab) and a section treated without antibody (-Ab). **C.** Representative signal of the LM20 (labelling methylesterified homogalacturonans, JIM5 and LM19 (labelling partially demethylesterified homogalacturonans) in WT and in *ap2-5*, *ap2-6*, *ap2-7* and *ap2-12* (crossed with WT pollen) mutant seeds at 3 and 6 DAP. Scale bars, 10 µm; n = 14 – 18 seeds, two independent experiments but JIM5 at 3DAP (n = 5 – 6 seeds, one experiment). **D.** Relative levels of the oligosaccharides released following digestion of the xyloglucans of intact seeds at 3, 6 and 9 DAP with an endocellulase. 2 independent experiments with 3 technical repeats, WT and *ap2-6* were compared using bilateral Student’s tests.

**Figure S5.**
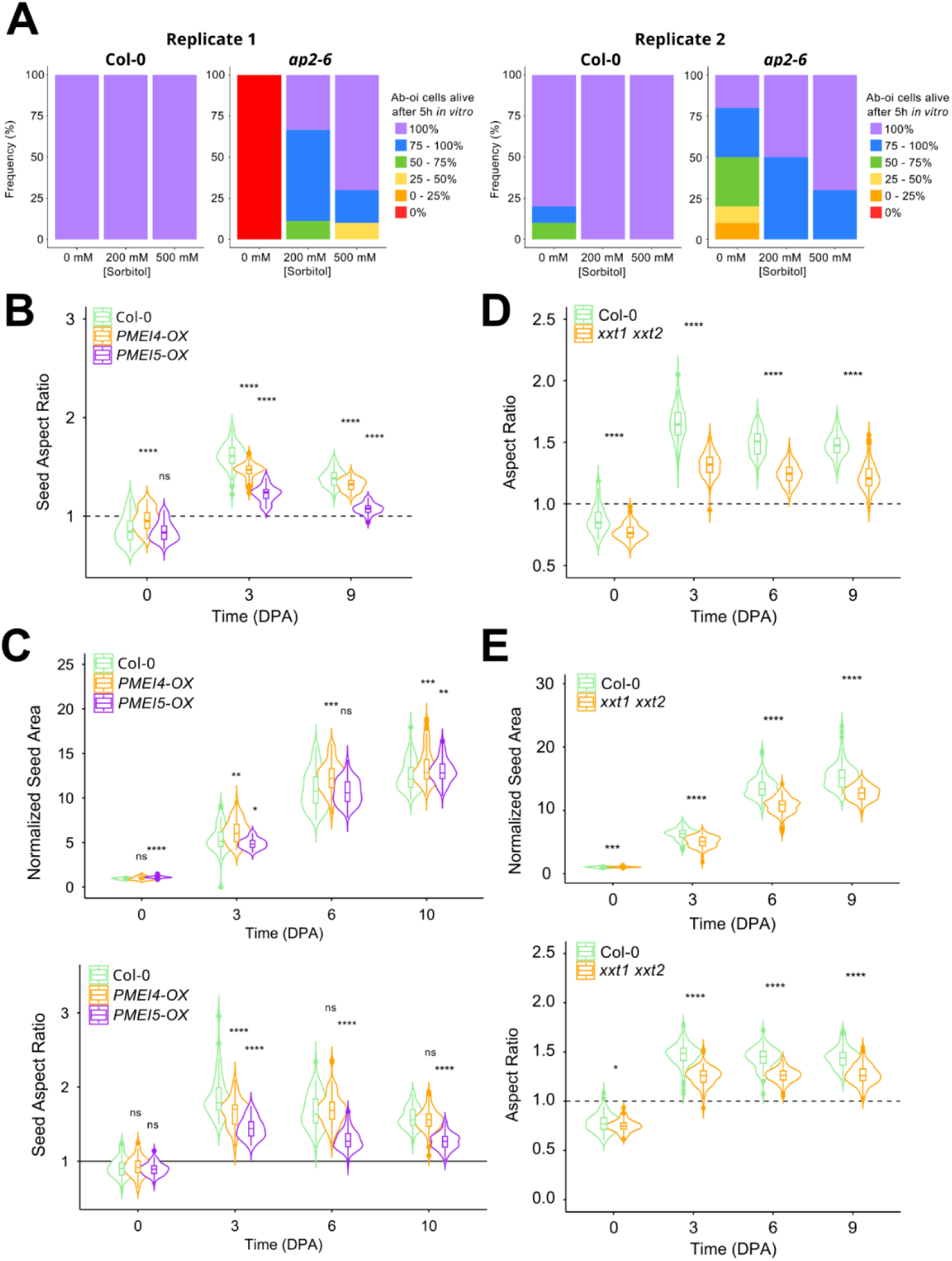
AP2-dependent changes in wall composition affect cell integrity and growth. **A.** Effect of medium osmolarity on ab-oi cell survival in WT and *ap2-6* seeds expressing a membrane marker at 2 DAP and cultivated *in vitro* for 5h; Replicate 1, *LTi6b-GFP* reporter, n = 8 – 10 seeds per condition; Replicate 2, *LTi6b-tdTomato* reporter, n = 10 seeds per condition. **B.** Analysis of the shape of WT, *PMEI4-OX* and *PMEI5-OX* seeds at 0, 3 and 9 DAP; n = 31 – 84 seeds from 3 siliques per day per genotype, one experiment. **C.** Growth dynamics of WT, *PMEI4-OX* and *PMEI5-OX* seeds at 0, 3, 6 and 9 DAP; n = 33 - 110 seeds from 3-4 siliques per day per genotype from one experiment independent from the one presented in Figure 5 and Figure S5B. Data were compared using bilateral Student’s tests. **D.** Analysis of the shape of WT and *xxt1 xxt2* seeds at 0, 3, 6 and 9 DAP; n = 42 – 124 seeds from 3 siliques per day per genotype, one experiment. Data were compared using bilateral Student’s tests. **E.** Growth dynamics of WT and *xxt1 xxt2* seeds at 0, 3, 6 and 9 DAP; n = 65 - 88 seeds from 3-4 siliques per day per genotype from one experiment independent from the one presented in Figure 5 and Figure S5D. Data were compared using bilateral Student’s tests.

**Figure S6:**
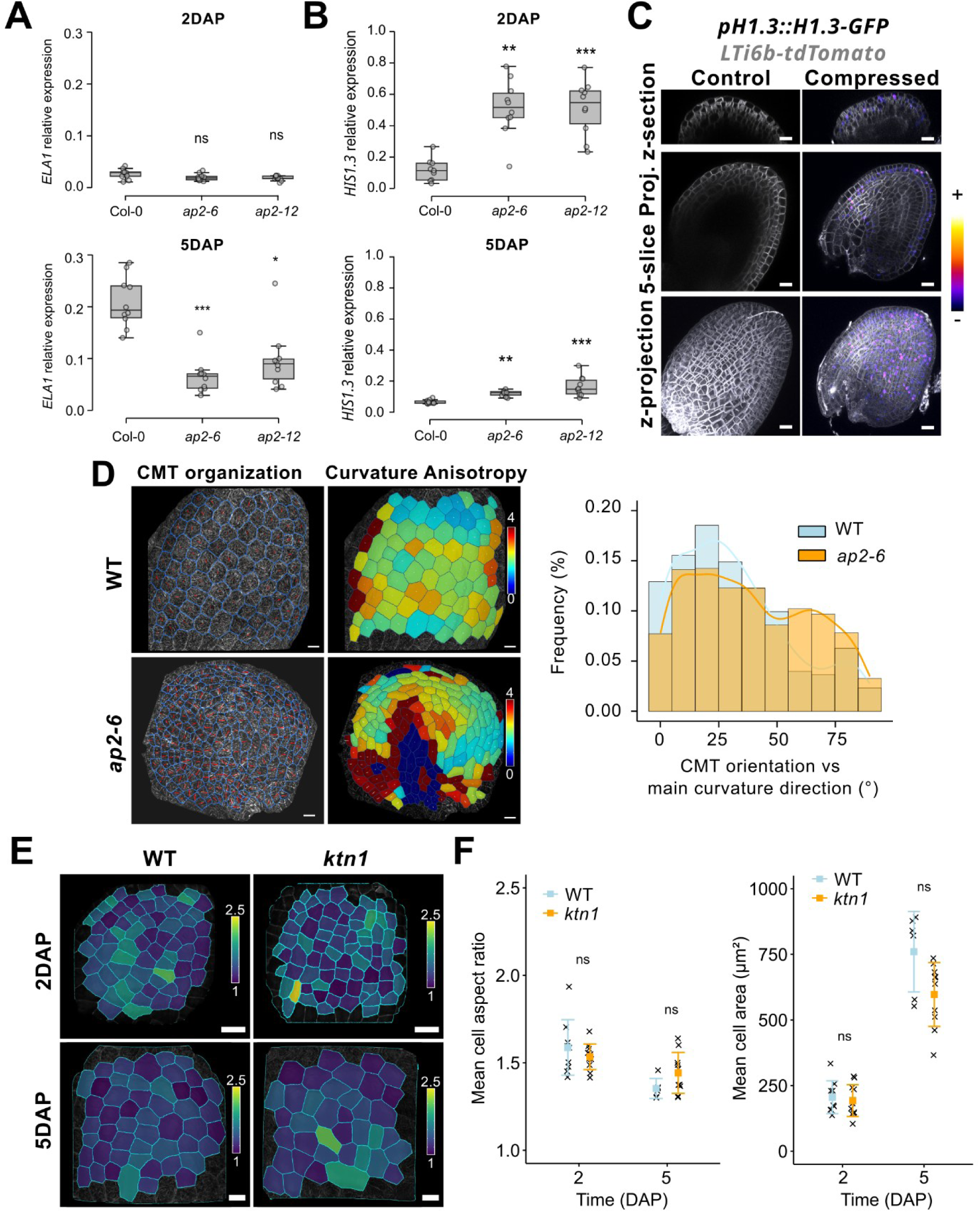
Mechanosensitive responses and shape determination. **A-B**. Relative expression of *ELA1* (B) and *H1.3* at 2 and 5 DAP in the WT and in the *ap2-6* and *ap2-12* mutants (crossed with WT pollen); n = 10 replicates from 2 independent experiments. Statistical differences established using the Kruskal-Wallis test followed by Dunn’s Post-hoc test. **C.** Representative pictures showing the expression of *H1.*3 assessed using the *pH1.3::H1.3-GFP* reporter in WT seeds after a 24h compression of the fruit using a microvice. **D.** Representative pictures and quantification of the correlation between the orientation of the CMTs (imaged using the *p35S::MAP65-1-RFP* reporter) facing the outer-side of the ab-oi layer and the curvature anisotropy in WT and *ap2-6* (crossed with WT pollen) seeds at 5DPA. Scale bars: 20 µm, n = 80 – 224 cells from 3 – 6 seeds per genotype, one experiment. For the CMT picture, each cell is highlighted in blue and the orientation of the red bars shows the mean orientation of the CMTs in each cell and their length, their degree of organization. In the heatmap of curvature anisotropy, the orientations of the two perpendicular white bars represent the axes of maximum and minimum curvature in each cell, and their lengths the degree of curvature in each of these two directions. **E.** Heatmap representation and quantification of the shape (aspect ratio) and size (area) of the cells in the ab-oi layer of WT and *ktn1* seeds at 2 and 5 DAP. Scale bars: 20 µm, n = 19 – 107 cells from 6 to 11 seeds, 2 independent experiments. Data were compared using bilateral Student’s tests.

## Supplementary Tables

**<u>Table S1:</u>** List of the differentially expressed genes (DEGs) in Col-0 *vs ap2-6* at 2 or 5 DAP with p-value < 0.05 obtained by bulk RNA sequencing (RNAseq).

**<u>Table S2:</u>** Expression of the genes normally enriched in the ab-oi layer (oi2) at 3 DAP from Martin *et al*, 2025^1^ (|log₂FC| ≥ 1, *p* ≤ 0.05) in WT and *ap2-6* seeds at 2 and 5 DAP (pct1, percent of nuclei expressing the gene in the cluster, pct2, percent of all other cell expressing the gene).

**<u>Table S3:</u>** Expression of the genes normally enriched in the ab-oi layer (oi2) at 5 DAP from Martin *et al*, 2025^1^ (|log₂FC| ≥ 1, *p* ≤ 0.05) in WT and *ap2-6* seeds at 2 and 5 DAP (pct1, percent of nuclei expressing the gene in the cluster, pct2, percent of all other cell expressing the gene).

**<u>Table S4:</u>** Expression of the genes normally enriched in the ad-oi layer (oi1) at 3 DAP from Martin *et al*, 2025^1^ (|log₂FC| ≥ 1, *p* ≤ 0.05) in WT and *ap2-6* seeds at 2 and 5 DAP (pct1, percent of nuclei expressing the gene in the cluster, pct2, percent of all other cell expressing the gene).

**<u>Table S5:</u>** Expression of the genes normally enriched in the ad-oi layer (oi1) at 5 DAP from Martin *et al*, 2025^1^ (|log₂FC| ≥ 1, *p* ≤ 0.05) in WT and *ap2-6* seeds at 2 and 5 DAP. (pct1, percent of nuclei expressing the gene in the cluster, pct2, percent of all other cell expressing the gene).

**<u>Table S6:</u>** List of putative AP2 direct targets extracted by Bertran Garcia de Olalla, E., Cerise, M., Rodríguez-Maroto, G. et al. *, 2024*^7^ from the ChIPseq (Chromatin Immunoprecipitation Sequencing) dataset of inflorescences of Yant *et al*, 2010^3^.

**<u>Table S7:</u>** Expression of the genes normally enriched in the ad-oi layer (oi1) at 3 DAP with a |log₂FC| ≥ 2, *p* ≤ 0.05 and a filtering for the keyword “transcription” from Martin *et al*, 2025^1^ in WT and *ap2-6* seeds at 2 and 5 DAP (pct1, percent of nuclei expressing the gene in the cluster, pct2, percent of all other cell expressing the gene).

**<u>Table S8:</u>** Expression of the genes normally enriched in the ad-oi layer (oi1) at 5 DAP with a |log₂FC| ≥ 2, *p* ≤ 0.05 and a filtering for the keyword “transcription” from Martin *et al*, 2025^1^ in WT and *ap2-6* seeds at 2 and 5 DAP (pct1, percent of nuclei expressing the gene in the cluster, pct2, percent of all other cell expressing the gene).

**<u>Table S9:</u>** Expression of the genes normally enriched in the ab-oi layer (oi2) at 3 DAP with a |log₂FC| ≥ 2, *p* ≤ 0.05 and a filtering for the keyword “transcription” from Martin *et al*, 2025^1^ in WT and *ap2-6* seeds at 2 and 5 DAP (pct1, percent of nuclei expressing the gene in the cluster, pct2, percent of all other cell expressing the gene).

**<u>Table S10:</u>** Expression of the genes normally enriched in the ab-oi layer (oi2) at 5 DAP with a |log₂FC| ≥ 2, *p* ≤ 0.05 and a filtering for the keyword “transcription” from Martin *et al*, 2025^1^ in WT and *ap2-6* seeds at 2 and 5 DAP (pct1, percent of nuclei expressing the gene in the cluster, pct2, percent of all other cell expressing the gene).

**<u>Table S11:</u>** Expression of the genes normally enriched in the ab-oi layer (oi2) at 3 DAP with a |log₂FC| ≥ 2, *p* ≤ 0.05 and a filtering for the keyword “wall” from Martin *et al*, 2025^1^ in WT and *ap2-6* seeds at 2 and 5 DAP (pct1, percent of nuclei expressing the gene in the cluster, pct2, percent of all other cell expressing the gene).

**<u>Table S12:</u>** Expression of the genes normally enriched in the ab-oi layer (oi2) at 5 DAP with a |log₂FC| ≥ 2, *p* ≤ 0.05 and a filtering for the keyword “wall” from Martin *et al*, 2025^1^ in WT and *ap2-6* seeds at 2 and 5 DAP (pct1, percent of nuclei expressing the gene in the cluster, pct2, percent of all other cell expressing the gene).

**<u>Table S13:</u>** Expression of the genes normally enriched in the ad-oi layer (oi1) at 3 DAP with a |log₂FC| ≥ 2, *p* ≤ 0.05 and a filtering for the keyword “wall” from Martin *et al*, 2025^1^ in WT and *ap2-6* seeds at 2 and 5 DAP (pct1, percent of nuclei expressing the gene in the cluster, pct2, percent of all other cell expressing the gene).

**<u>Table S14:</u>** Expression of the genes normally enriched in the ad-oi layer (oi1) at 5 DAP with a |log₂FC| ≥ 2, *p* ≤ 0.05 and a filtering for the keyword “wall” from Martin *et al*, 2025^1^ in WT and *ap2-6* seeds at 2 and 5 DAP (pct1, percent of nuclei expressing the gene in the cluster, pct2, percent of all other cell expressing the gene).

**<u>Table S15:</u>** Description of the mutants and transgenic lines used in this study

**<u>Table S16:</u>** Sequence of the primers used in this study

## References

1. Landrein, B., and Ingram, G. (2019). Connected through the force: Mechanical signals in plant development. J. Exp. Bot. 10.1093/jxb/erz103.

2. Kutschera, U., and Niklas, K.J. (2007). The epidermal-growth-control theory of stem elongation: An old and a new perspective. Journal of Plant Physiology 164, 1395–1409. 10.1016/j.jplph.2007.08.002.

3. Kelly-Bellow, R., Lee, K., Kennaway, R., Barclay, J.E., Whibley, A., Bushell, C., Spooner, J., Yu, M., Brett, P., Kular, B., et al. (2023). Brassinosteroid coordinates cell layer interactions in plants via cell wall and tissue mechanics. Science 380, 1275–1281. 10.1126/science.adf0752.

4. Savaldi-Goldstein, S., Peto, C., and Chory, J. (2007). The epidermis both drives and restricts plant shoot growth. Nature 446, 199–202. 10.1038/nature05618.

5. Kierzkowski, D., Nakayama, N., Routier-Kierzkowska, A.-L., Weber, A., Bayer, E., Schorderet, M., Reinhardt, D., Kuhlemeier, C., and Smith, R.S. (2012). Elastic Domains Regulate Growth and Organogenesis in the Plant Shoot Apical Meristem. Science 335, 1096–1099. 10.1126/science.1213100.

6. Landrein, B., Kiss, A., Sassi, M., Chauvet, A., Das, P., Cortizo, M., Laufs, P., Takeda, S., Aida, M., Traas, J., et al. (2015). Mechanical stress contributes to the expression of the STM homeobox gene in Arabidopsis shoot meristems. Elife 4, e07811. 10.7554/eLife.07811.

7. Hamant, O., Heisler, M.G., Jonsson, H., Krupinski, P., Uyttewaal, M., Bokov, P., Corson, F., Sahlin, P., Boudaoud, A., Meyerowitz, E.M., et al. (2008). Developmental Patterning by Mechanical Signals in Arabidopsis. Science 322, 1650–1655. 10.1126/science.1165594.

8. Sampathkumar, A., Krupinski, P., Wightman, R., Milani, P., Berquand, A., Boudaoud, A., Hamant, O., Jönsson, H., and Meyerowitz, E.M. (2014). Subcellular and supracellular mechanical stress prescribes cytoskeleton behavior in Arabidopsis cotyledon pavement cells. eLife 3, e01967. 10.7554/eLife.01967.

9. Heisler, M.G., Hamant, O., Krupinski, P., Uyttewaal, M., Ohno, C., Jönsson, H., Traas, J., and Meyerowitz, E.M. (2010). Alignment between PIN1 Polarity and Microtubule Orientation in the Shoot Apical Meristem Reveals a Tight Coupling between Morphogenesis and Auxin Transport. PLoS Biology 8, e1000516. 10.1371/journal.pbio.1000516.

10. Iida, H., Mähönen, A.P., Jürgens, G., and Takada, S. (2023). Epidermal injury-induced derepression of key regulator ATML1 in newly exposed cells elicits epidermis regeneration. Nat Commun 14, 1031. 10.1038/s41467-023-36731-6.

11. Johnson, K.L., Degnan, K.A., Ross Walker, J., and Ingram, G.C. (2005). AtDEK1 is essential for specification of embryonic epidermal cell fate: AtDEK1 in epidermal cell fate. The Plant Journal 44, 114–127. 10.1111/j.1365-313X.2005.02514.x.

12. Ogawa, E., Yamada, Y., Sezaki, N., Kosaka, S., Kondo, H., Kamata, N., Abe, M., Komeda, Y., and Takahashi, T. (2015). ATML1 and PDF2 Play a Redundant and Essential Role in Arabidopsis Embryo Development. Plant and Cell Physiology 56, 1183–1192. 10.1093/pcp/pcv045.

13. Ohto, M., Stone, S.L., and Harada, J.J. (2007). Genetic Control of Seed Development and Seed Mass. In Seed Development, Dormancy and Germination, K. J. Bradford and H. Nonogaki, eds. (Blackwell Publishing Ltd), pp. 1–24. 10.1002/9780470988848.ch1.

14. Braat, J., and Landrein, B. (2025). Mechanical control of plant organ growth: Lessons from the seed. Current Opinion in Plant Biology 85, 102737. 10.1016/j.pbi.2025.102737.

15. Creff, A., Brocard, L., and Ingram, G. (2015). A mechanically sensitive cell layer regulates the physical properties of the Arabidopsis seed coat. Nat Commun 6, 6382. 10.1038/ncomms7382.

16. Creff, A., Ali, O., Bied, C., Bayle, V., Ingram, G., and Landrein, B. (2023). Evidence that endosperm turgor pressure both promotes and restricts seed growth and size. Nat Commun 14, 67. 10.1038/s41467-022-35542-5.

17. Bauer, A., Ali, O., Bied, C., Bœuf, S., Bovio, S., Delattre, A., Ingram, G., Golz, J.F., and Landrein, B. (2024). Spatiotemporally distinct responses to mechanical forces shape the developing seed of Arabidopsis. EMBO J 43, 2733–2758. 10.1038/s44318-024-00138-w.

18. Brambilla, V., Kater, M., and Colombo, L. (2008). Ovule integument identity determination in Arabidopsis. Plant Signaling & Behavior 3, 246–247. 10.4161/psb.3.4.5175.

19. Bowman, J.L., and Moyroud, E. (2024). Reflections on the ABC model of flower development. The Plant Cell 36, 1334–1357. 10.1093/plcell/koae044.

20. Jofuku, K.D., Omidyar, P.K., Gee, Z., and Okamuro, J.K. (2005). Control of seed mass and seed yield by the floral homeotic gene APETALA2. Proc. Natl. Acad. Sci. U.S.A. 102, 3117–3122. 10.1073/pnas.0409893102.

21. Ohto, M., Floyd, S.K., Fischer, R.L., Goldberg, R.B., and Harada, J.J. (2009). Effects of APETALA2 on embryo, endosperm, and seed coat development determine seed size in Arabidopsis. Sex Plant Reprod 22, 277–289. 10.1007/s00497-009-0116-1.

22. Wakem, M.P. (2003). Mutation in the ap2-6 allele causes recognition of a cryptic splice site. Journal of Experimental Botany 54, 2655–2660. 10.1093/jxb/erg298.

23. Dean, G., Cao, Y., Xiang, D., Provart, N.J., Ramsay, L., Ahad, A., White, R., Selvaraj, G., Datla, R., and Haughn, G. (2011). Analysis of Gene Expression Patterns during Seed Coat Development in Arabidopsis. Molecular Plant 4, 1074–1091. 10.1093/mp/ssr040.

24. Bertran Garcia De Olalla, E., Cerise, M., Rodríguez-Maroto, G., Casanova-Ferrer, P., Vayssières, A., Severing, E., López Sampere, Y., Wang, K., Schäfer, S., Formosa-Jordan, P., et al. (2024). Coordination of shoot apical meristem shape and identity by APETALA2 during floral transition in Arabidopsis. Nat Commun 15, 6930. 10.1038/s41467-024-51341-6.

25. Yant, L., Mathieu, J., Dinh, T.T., Ott, F., Lanz, C., Wollmann, H., Chen, X., and Schmid, M. (2010). Orchestration of the Floral Transition and Floral Development in *Arabidopsis* by the Bifunctional Transcription Factor APETALA2. The Plant Cell 22, 2156–2170. 10.1105/tpc.110.075606.

26. Kunst, L., Klenz, J.E., Martinez-Zapater, J., and Haughn, G.W. (1989). AP2 Gene Determines the Identity of Perianth Organs in Flowers of Arabidopsis thaliana. Plant Cell, 1195–1208. 10.1105/tpc.1.12.1195.

27. Lima, R.B., Pankaj, R., Ehlert, S.T., Finger, P., Fröhlich, A., Bayle, V., Landrein, B., Sampathkumar, A., and Figueiredo, D.D. (2024). Seed coat-derived brassinosteroid signaling regulates endosperm development. Nat Commun 15, 9352. 10.1038/s41467-024-53671-x.

28. Rolletschek, H., Muszynska, A., Schwender, J., Radchuk, V., Heinemann, B., Hilo, A., Plutenko, I., Keil, P., Ortleb, S., Wagner, S., et al. (2024). Mechanical forces orchestrate the metabolism of the developing oilseed rape embryo. New Phytologist 244, 1328–1344. 10.1111/nph.19990.

29. Belmonte, M.F., Kirkbride, R.C., Stone, S.L., Pelletier, J.M., Bui, A.Q., Yeung, E.C., Hashimoto, M., Fei, J., Harada, C.M., Munoz, M.D., et al. (2013). Comprehensive developmental profiles of gene activity in regions and subregions of the Arabidopsis seed. Proceedings of the National Academy of Sciences 110, E435–E444. 10.1073/pnas.1222061110.

30. Martin, C.A., Cogdill, K.R., Pusey, A.L., and Gehring, M. (2025). A transcriptional atlas of early Arabidopsis seed development suggests mechanisms for inter-tissue coordination. Preprint at Plant Biology, 10.1101/2025.04.29.651360 https://doi.org/10.1101/2025.04.29.651360.

31. Picard, C.L., Povilus, R.A., Williams, B.P., and Gehring, M. (2021). Transcriptional and imprinting complexity in Arabidopsis seeds at single-nucleus resolution. Nat. Plants 7, 730–738. 10.1038/s41477-021-00922-0.

32. Schrick, K., Ahmad, B., and Nguyen, H.V. (2023). HD-Zip IV transcription factors: Drivers of epidermal cell fate integrate metabolic signals. Current Opinion in Plant Biology 75, 102417. 10.1016/j.pbi.2023.102417.

33. Abe, M., Katsumata, H., Komeda, Y., and Takahashi, T. (2003). Regulation of shoot epidermal cell differentiation by a pair of homeodomain proteins in *Arabidopsis*. Development 130, 635–643. 10.1242/dev.00292.

34. Szymanski, D.B., Jilk, R.A., Pollock, S.M., and Marks, M.D. (1998). Control of *GL2* expression in *Arabidopsis* leaves and trichomes. Development 125, 1161–1171. 10.1242/dev.125.7.1161.

35. Abe, M., Takahashi, T., and Komeda, Y. (1999). Cloning and Characterization of an L1 Layer-Specific Gene in Arabidopsis thaliana. Plant and Cell Physiology 40, 571–580. 10.1093/oxfordjournals.pcp.a029579.

36. Golz, J.F., Allen, P.J., Li, S.F., Parish, R.W., Jayawardana, N.U., Bacic, A., and Doblin, M.S. (2018). Layers of regulation – Insights into the role of transcription factors controlling mucilage production in the Arabidopsis seed coat. Plant Science 272, 179–192. 10.1016/j.plantsci.2018.04.021.

37. Allen, P.J., Napoli, R.S., Parish, R.W., and Li, S.F. (2023). MYB-bHLH-TTG1 in a Multi-tiered Pathway Regulates *Arabidopsis* Seed Coat Mucilage Biosynthesis Genes Including *PECTIN METHYLESTERASE INHIBITOR14* Required for Homogalacturonan Demethylesterification. Plant And Cell Physiology 64, 906–919. 10.1093/pcp/pcad065.

38. Dubos, C., Le Gourrierec, J., Baudry, A., Huep, G., Lanet, E., Debeaujon, I., Routaboul, J., Alboresi, A., Weisshaar, B., and Lepiniec, L. (2008). MYBL2 is a new regulator of flavonoid biosynthesis in *Arabidopsis thaliana*. The Plant Journal 55, 940–953. 10.1111/j.1365-313X.2008.03564.x.

39. Jiang, W., Yin, Q., Liu, J., Su, X., Han, X., Li, Q., Zhang, J., and Pang, Y. (2024). The APETALA2–MYBL2 module represses proanthocyanidin biosynthesis by affecting formation of the MBW complex in seeds of Arabidopsis thaliana. Plant Communications 5, 100777. 10.1016/j.xplc.2023.100777.

40. Broun, P. (2005). Transcriptional control of flavonoid biosynthesis: a complex network of conserved regulators involved in multiple aspects of differentiation in Arabidopsis. Current Opinion in Plant Biology 8, 272–279. 10.1016/j.pbi.2005.03.006.

41. Johnson, C.S., Kolevski, B., and Smyth, D.R. (2002). *TRANSPARENT TESTA GLABRA2*, a Trichome and Seed Coat Development Gene of Arabidopsis, Encodes a WRKY Transcription Factor. Plant Cell 14, 1359–1375. 10.1105/tpc.001404.

42. Garcia, D., Fitz Gerald, J.N., and Berger, F. (2005). Maternal control of integument cell elongation and zygotic control of endosperm growth are coordinated to determine seed size in Arabidopsis. Plant Cell 17, 52–60. 10.1105/tpc.104.027136.

43. Cosgrove, D.J. (2022). Building an extensible cell wall. Plant Physiology 189, 1246–1277. 10.1093/plphys/kiac184.

44. Hocq, L., Pelloux, J., and Lefebvre, V. (2017). Connecting Homogalacturonan-Type Pectin Remodeling to Acid Growth. Trends Plant Sci 22, 20–29. 10.1016/j.tplants.2016.10.009.

45. Marcus, S.E., Verhertbruggen, Y., Hervé, C., Ordaz-Ortiz, J.J., Farkas, V., Pedersen, H.L., Willats, W.G., and Knox, J.P. (2008). Pectic homogalacturonan masks abundant sets of xyloglucan epitopes in plant cell walls. BMC Plant Biol 8, 60. 10.1186/1471-2229-8-60.

46. Pelletier, S., Van Orden, J., Wolf, S., Vissenberg, K., Delacourt, J., Ndong, Y.A., Pelloux, J., Bischoff, V., Urbain, A., Mouille, G., et al. (2010). A role for pectin de-methylesterification in a developmentally regulated growth acceleration in dark-grown Arabidopsis hypocotyls. New Phytologist 188, 726–739. 10.1111/j.1469-8137.2010.03409.x.

47. Wolf, S., Mravec, J., Greiner, S., Mouille, G., and Höfte, H. (2012). Plant cell wall homeostasis is mediated by brassinosteroid feedback signaling. Curr. Biol. 22, 1732–1737. 10.1016/j.cub.2012.07.036.

48. Zabotina, O.A., Avci, U., Cavalier, D., Pattathil, S., Chou, Y.-H., Eberhard, S., Danhof, L., Keegstra, K., and Hahn, M.G. (2012). Mutations in Multiple *XXT* Genes of Arabidopsis Reveal the Complexity of Xyloglucan Biosynthesis. Plant Physiology 159, 1367–1384. 10.1104/pp.112.198119.

49. Cosgrove, D.J. (2024). Structure and growth of plant cell walls. Nat Rev Mol Cell Biol 25, 340–358. 10.1038/s41580-023-00691-y.

50. Zhang, Y., Yu, J., Wang, X., Durachko, D.M., Zhang, S., and Cosgrove, D.J. (2021). Molecular insights into the complex mechanics of plant epidermal cell walls. Science 372, 706–711. 10.1126/science.abf2824.

51. Fal, K., Korsbo, N., Alonso-Serra, J., Teles, J., Liu, M., Refahi, Y., Chabouté, M.-E., Jönsson, H., and Hamant, O. (2021). Tissue folding at the organ–meristem boundary results in nuclear compression and chromatin compaction. PNAS 118. 10.1073/pnas.2017859118.

52. Rutowicz, K., Puzio, M., Halibart-Puzio, J., Lirski, M., Kroteń, M.A., Kotliński, M., Kniżewski, Ł., Lange, B., Muszewska, A., Śniegowska-Świerk, K., et al. (2015). A specialized histone H1 variant is required for adaptive responses to complex abiotic stress and related DNA methylation in Arabidopsis. Plant Physiol., pp.00493.2015. 10.1104/pp.15.00493.

53. Alonso-Serra, J., Cheddadi, I., Kiss, A., Cerutti, G., Lang, M., Dieudonné, S., Lionnet, C., Godin, C., and Hamant, O. (2024). Water fluxes pattern growth and identity in shoot meristems. Nat Commun 15, 6944. 10.1038/s41467-024-51099-x.

54. Dauphin, B.G., Ropartz, D., Ranocha, P., Rouffle, M., Carton, C., Le Ru, A., Martinez, Y., Fourquaux, I., Ollivier, S., Mac-Bear, J., et al. (2024). TBL38 atypical homogalacturonan-acetylesterase activity and cell wall microdomain localization in Arabidopsis seed mucilage secretory cells. iScience 27, 109666. 10.1016/j.isci.2024.109666.

55. Philippe, F., Pelloux, J., and Rayon, C. (2017). Plant pectin acetylesterase structure and function: new insights from bioinformatic analysis. BMC Genomics 18, 456. 10.1186/s12864-017-3833-0.

56. Shahin, L., Zhang, L., Mohnen, D., and Urbanowicz, B.R. (2023). Insights into pectin O-acetylation in the plant cell wall: structure, synthesis, and modification. The Cell Surface 9, 100099. 10.1016/j.tcsw.2023.100099.

57. Di Marzo, M., Babolin, N., Viana, V.E., De Oliveira, A.C., Gugi, B., Caporali, E., Herrera-Ubaldo, H., Martínez-Estrada, E., Driouich, A., De Folter, S., et al. (2022). The Genetic Control of SEEDSTICK and LEUNIG-HOMOLOG in Seed and Fruit Development: New Insights into Cell Wall Control. Plants 11, 3146. 10.3390/plants11223146.

58. Ezquer, I., Mizzotti, C., Nguema-Ona, E., Gotté, M., Beauzamy, L., Viana, V.E., Dubrulle, N., Costa De Oliveira, A., Caporali, E., Koroney, A.-S., et al. (2016). The Developmental Regulator SEEDSTICK Controls Structural and Mechanical Properties of the Arabidopsis Seed Coat. Plant Cell 28, 2478–2492. 10.1105/tpc.16.00454.

59. Di Marzo, M., Viana, V.E., Banfi, C., Cassina, V., Corti, R., Herrera-Ubaldo, H., Babolin, N., Guazzotti, A., Kiegle, E., Gregis, V., et al. (2022). Cell wall modifications by α-XYLOSIDASE1 are required for control of seed and fruit size in Arabidopsis. Journal of Experimental Botany 73, 1499–1515. 10.1093/jxb/erab514.

60. Yu, B., He, X., Tang, Y., Chen, Z., Zhou, L., Li, X., Zhang, C., Huang, X., Yang, Y., Zhang, W., et al. (2023). Photoperiod controls plant seed size in a CONSTANS-dependent manner. Nat. Plants. 10.1038/s41477-023-01350-y.

61. Peaucelle, A., Wightman, R., and Höfte, H. (2015). The Control of Growth Symmetry Breaking in the Arabidopsis Hypocotyl. Current Biology 25, 1746–1752. 10.1016/j.cub.2015.05.022.

62. Figueiredo, D.D., Batista, R.A., Roszak, P.J., Hennig, L., and Köhler, C. (2016). Auxin production in the endosperm drives seed coat development in *Arabidopsis*. eLife 5, e20542. 10.7554/eLife.20542.

63. Pankaj, R., Lima, R.B., Luo, G.-Y., Ehlert, S., Del Toro-de León, G., Bente, H., Finger, P., Sato, H., and Figueiredo, D.D. (2023). BRI1-mediated removal of seed coat H3K27me3 marks is a brassinosteroid-independent process. Preprint, 10.1101/2023.12.07.569203 https://doi.org/10.1101/2023.12.07.569203.

64. Gonzalez, A., Mendenhall, J., Huo, Y., and Lloyd, A. (2009). TTG1 complex MYBs, MYB5 and TT2, control outer seed coat differentiation. Developmental Biology 325, 412–421. 10.1016/j.ydbio.2008.10.005.

65. Nemec-Venza, Z., Kiss, A., German, N., Bovio, S., Field, N., Martin, M., Lionnet, C., Vavrdova, T., Feng, Q., De Winter, F., et al. (2025). Plant cells at the organ surface use mechanical cues to activate a specific growth control programme. Preprint at Plant Biology, 10.1101/2025.08.25.672229 https://doi.org/10.1101/2025.08.25.672229.

66. Clough, S.J., and Bent, A.F. (1998). Floral dip: a simplified method for Agrobacterium - mediated transformation of Arabidopsis thaliana. The Plant Journal 16, 735–743. 10.1046/j.1365-313x.1998.00343.x.

67. Lambert, I., Paysant-Le Roux, C., Colella, S., and Martin-Magniette, M.-L. (2020). DiCoExpress: a tool to process multifactorial RNAseq experiments from quality controls to co-expression analysis through differential analysis based on contrasts inside GLM models. Plant Methods 16, 68. 10.1186/s13007-020-00611-7.

68. Besten, M., Hendriksz, M., Michels, L., Charrier, B., Smakowska-Luzan, E., Weijers, D., Borst, J.W., and Sprakel, J. (2024). CarboTag: a modular approach for live and functional imaging of plant cell walls. Preprint, 10.1101/2024.07.05.597952 https://doi.org/10.1101/2024.07.05.597952.

69. Boudaoud, A., Burian, A., Borowska-Wykręt, D., Uyttewaal, M., Wrzalik, R., Kwiatkowska, D., and Hamant, O. (2014). FibrilTool, an ImageJ plug-in to quantify fibrillar structures in raw microscopy images. Nat Protoc 9, 457–463. 10.1038/nprot.2014.024.

70. Prabhullachandran, U., Urbánková, I., Medaglia-Mata, A., Creff, A., Voxeur, A., Petřík, I., Pěnčík, A., Novák, O., Landrein, B., Hejátko, J., et al. (2024). Long-term high temperatures affect seed maturation and seed coat integrity in *Brassica napus*. Preprint, 10.1101/2024.11.27.625589 https://doi.org/10.1101/2024.11.27.625589.

71. Voxeur, A., Habrylo, O., Guénin, S., Miart, F., Soulié, M.-C., Rihouey, C., Pau-Roblot, C., Domon, J.-M., Gutierrez, L., Pelloux, J., et al. (2019). Oligogalacturonide production upon *Arabidopsis thaliana* – *Botrytis cinerea* interaction. Proc. Natl. Acad. Sci. U.S.A. 116, 19743–19752. 10.1073/pnas.1900317116.

## Supplementary References

1. Martin, C.A., Cogdill, K.R., Pusey, A.L., and Gehring, M. (2025). A transcriptional atlas of early Arabidopsis seed development suggests mechanisms for inter-tissue coordination. Preprint at Plant Biology, 10.1101/2025.04.29.651360 https://doi.org/10.1101/2025.04.29.651360.

2. Belmonte, M.F., Kirkbride, R.C., Stone, S.L., Pelletier, J.M., Bui, A.Q., Yeung, E.C., Hashimoto, M., Fei, J., Harada, C.M., Munoz, M.D., et al. (2013). Comprehensive developmental profiles of gene activity in regions and subregions of the Arabidopsis seed. Proceedings of the National Academy of Sciences 110, E435–E444. 10.1073/pnas.1222061110.

3. Yant, L., Mathieu, J., Dinh, T.T., Ott, F., Lanz, C., Wollmann, H., Chen, X., and Schmid, M. (2010). Orchestration of the Floral Transition and Floral Development in *Arabidopsis* by the Bifunctional Transcription Factor APETALA2. The Plant Cell 22, 2156–2170. 10.1105/tpc.110.075606.

4. Golz, J.F., Allen, P.J., Li, S.F., Parish, R.W., Jayawardana, N.U., Bacic, A., and Doblin, M.S. (2018). Layers of regulation – Insights into the role of transcription factors controlling mucilage production in the Arabidopsis seed coat. Plant Science 272, 179–192. 10.1016/j.plantsci.2018.04.021.

5. Dubos, C., Le Gourrierec, J., Baudry, A., Huep, G., Lanet, E., Debeaujon, I., Routaboul, J., Alboresi, A., Weisshaar, B., and Lepiniec, L. (2008). MYBL2 is a new regulator of flavonoid biosynthesis in *Arabidopsis thaliana*. The Plant Journal 55, 940–953. 10.1111/j.1365-313X.2008.03564.x.

6. Jiang, W., Yin, Q., Liu, J., Su, X., Han, X., Li, Q., Zhang, J., and Pang, Y. (2024). The APETALA2–MYBL2 module represses proanthocyanidin biosynthesis by affecting formation of the MBW complex in seeds of Arabidopsis thaliana. Plant Communications 5, 100777. 10.1016/j.xplc.2023.100777.

7. Bertran Garcia De Olalla, E., Cerise, M., Rodríguez-Maroto, G., Casanova-Ferrer, P., Vayssières, A., Severing, E., López Sampere, Y., Wang, K., Schäfer, S., Formosa-Jordan, P., et al. (2024). Coordination of shoot apical meristem shape and identity by APETALA2 during floral transition in Arabidopsis. Nat Commun 15, 6930. 10.1038/s41467-024-51341-6.

